# Minimizing inferential bias in the theory and design of nutritional experiments through the application of the equilateral mixture triangle

**DOI:** 10.64898/2025.12.06.692245

**Authors:** Colin Lynch, Kaitlin Baudier, Douglas Montgomery, Meghan Barrett

## Abstract

Animal nutritionists seek to understand how animals regulate the intake and balance of multiple nutrients, yet the design and analysis of such experiments are often limited by how nutrient spaces are represented. The geometric framework for nutrition (GFN) provides a powerful means to visualize nutrient interactions, but when more than two nutrients are considered, most empirical studies employ right-angled mixture triangles (RMTs). While convenient, this coordinate system can introduce visual artifacts that can bias inference. Here, we reintroduce the equilateral mixture triangle (EMT) as a complementary and biologically meaningful framework that preserves proportional relationships among nutrients and connects directly with the design principles of mixture experiments in engineering disciplines, where it is more commonly known as a simplex. The simplex provides a consistent bridge between theory and experimentation, allowing the same geometric representation to be used for choice and no-choice assays. For choice experiments, we extend GFN theory to show that animal feeding trajectories are bounded by the convex hull of available foods, show how it can scale to higher-dimensional nutrient systems, and develop statistical tests to distinguish between random choice and the defense of an intake target. For no-choice experiments, we demonstrate how simplex mixture designs can be integrated with response surface methodologies to model multivariate performance landscapes, identify optimal nutrient ratios, and avoid biologically unrealistic regions of nutrient space. For both types of experiments, we provide simulated case studies on how to design studies around the simplex. Together, these developments refine both the theoretical and experimental foundations of nutritional ecology, providing clear guidance for designing, visualizing, and interpreting multidimensional studies of nutrient regulation with diverse applications across ecological and agricultural disciplines.

## 1. Introduction

Nutrition is a cornerstone of biology, influencing processes that range from individual health or fitness to population dynamics, agricultural yield, and ecosystem stability [1]. Yet, nutrition is notoriously difficult to study, as foods are complex mixtures of interacting components whose availability is both dynamic and stochastic [2]. The geometric framework of nutrition (GFN) was developed to render this complexity tractable by representing animal feeding decisions and performance outcomes in a reduced-dimensional state-space [3]. The GFN provides a means to construct models that integrate diet, nutritional state, feeding behavior, physiology, and the consequences of nutrition across biological scales, from gene expression to life-history outcomes [4, 5, 6, 7]. The framework has now been applied to approximately fifty taxa, ranging from slime molds to invertebrates, fish, and numerous mammals, including humans [8]. It has also been used to address a wide variety of questions: how animals balance nutrient intake to achieve homeostasis, how physiological state influences diet choice, how nutrients interact to shape fitness landscapes, how to improve diets to increase yield, health, or well-being in domestic or farmed animals, and how these processes scale up to affect populations and ecosystems [9, 10, 11, 12, 13, 14].

While powerful, most applications of the GFN have been largely restricted to two nutritional axes (generally, carbohydrates and protein), limiting the ability to disentangle higher-order interactions among the three primary macronutrients: protein, carbohydrate, and lipid [15]. To extend beyond two dimensions, researchers introduced the right-angled mixture triangle (RMT) and the equilateral mixture triangle (EMT), both of which project three nutrient ratios into a two-dimensional plane [16]. Although most studies employing mixture triangles have focused on macronutrients, these coordinate systems can, in principle, represent any three-component system. For instance, both RMT and EMT formulations have been used to explore mixtures of different sugars [17, 16], and they could potentially be used to represent elemental ratios such as carbon, nitrogen, and phosphorus in stoichiometry [18]. Of the two, the RMT has been more widely adopted in field and laboratory studies [19, 20, 21, 22, 23, 24, 25, 26]. The primary argument for the RMT is that two of its axes are perpendicular, and are thus more interpretable to those accustomed to Cartesian coordinates [16]. However, to accomplish this, the method introduces distortions in the nutrient space by stretching the remaining axis, thereby magnifying the apparent influence of an arbitrarily chosen nutrient. These distortions can complicate both visualization and modeling. The EMT, in contrast, avoids this issue and has a long history of rigorous application in industrial engineering through mixture experiments, where it is known as a simplex (for definitions, see Table 1; [27, 28, 29]). This provides an opportunity to translate established methods for optimal experimental designs (Table 1 for explicit definition of ’optimal’ in this case) and hypothesis testing from the engineering tradition into nutritional ecology and animal sciences [30, 31].

**Table 1:** Key terms from mixture experiments and the geometric framework of nutrition.

| Term | Definition (as used here) |
| --- | --- |
| Simplex | Geometric set of all mixtures whose component proportions sum to a constant. For three components, the simplex is an equilateral triangle; for four, a tetrahedron. |
| Equilateral Mixture Triangle (EMT) | Ternary diagram for three-component mixtures drawn as an equilateral triangle. All directions are scaled equally, so distances and variation are not axis-distorted. |
| Right-Angled Mixture Triangle (RMT) | Ternary representation aligned with Cartesian axes, where two components are explicit and the third is implicit along a diagonal. It stretches space along the implicit axis and distorts distances relative to the EMT. |
| Mixture experiment / no-choice experiment | Experiment where each unit receives a single fixed composition (diet), and responses (e.g., growth, survival) are modeled as functions of that composition. |
| Choice experiment (choice assay) | Experiment where organisms choose among two or more foods with different compositions, generating trajectories through nutrient space that may converge on an intake target. |
| GFN (Geometric Framework of Nutrition) | Framework that represents nutrition in a multivariate state space, describing foods, intake targets, and feeding trajectories in terms of nutrient amounts or ratios. |
| Intake target | Nutrient ratio (and sometimes total amount) that animals regulate toward to maximize a focal suite of fitness-related traits. Appears as a point in nutrient space or in the simplex. |
| Growth target / nutrient target | Targets describing the composition of retained tissue (growth) or retained plus metabolized nutrients (nutrient). They may differ from the intake target and can lie outside the convex hull of available foods. |
| Food point | Point in the simplex representing the fixed nutrient ratio of a particular food or diet. |
| Convex hull / nutrient space polygon | Smallest convex polygon enclosing all food points. It represents the set of attainable nutritional states given the offered foods; trajectories cannot leave it without adding or changing foods. |
| Center of mass (mean composition) | Average composition of all offered foods under equal weighting. Under random choice (all foods equally likely), trajectories converge on this point. |
| Rule of compromise | Strategy for dealing with unbalanced or constrained diets (e.g., random choice, food-item bias, macronutrient bias, closest-distance optimization). Here, rules are expressed as probabilistic decision rules and trajectories in the simplex. |
| Closest-distance optimization | Rule of compromise where the animal repeatedly chooses the food that minimizes the next-step distance to the intake target in nutrient space, producing a piecewise path that approximates a straight line to the target. |
| Simplex-lattice design | Mixture design in which component proportions take discrete levels (e.g., $0, 1/k, \dots, 1$) subject to the sum constraint, forming an evenly spaced grid of compositions in the simplex. |
| Scheffé mixture model | Polynomial regression model for mixture data (linear, quadratic, special cubic, full cubic) that expresses the response as a function of component proportions and their products, while respecting $r_1 + r_2 + r_3 = 1$. |
| Alphabet-optimal design (A-, C-, D-optimal, etc.) | Design chosen to optimize a statistical efficiency criterion based on the information matrix (e.g., A-optimal: minimize average coefficient variance; D-optimal: maximize overall information). |
| Response surface | Predicted response over the simplex (or other designs) from a fitted model, showing how a response changes with composition. Peaks, ridges, and valleys indicate optima and trade-offs. |
| Desirability function | Transformation that maps each response to $[0, 1]$ ($0$ = unacceptable, $1$ = ideal), then combines them (e.g., by a geometric mean) into a single objective for multi-response optimization. |
| Pareto frontier (Pareto-optimal mixtures) | Set of mixtures for which no response can be improved without worsening at least one other. Represents trade-off-efficient nutrient ratios. |
| Mixture-process variable (MPV) design | Design that combines mixture components with one or more non-mixture (process) variables (e.g., temperature, micronutrient level), allowing interactions between composition and process conditions. |
| Constrained mixture region | Subset of the simplex defined by bounds on component proportions (e.g., welfare or palatability limits). Designs and optimality criteria are evaluated only within this feasible region. |

**Table 2:**
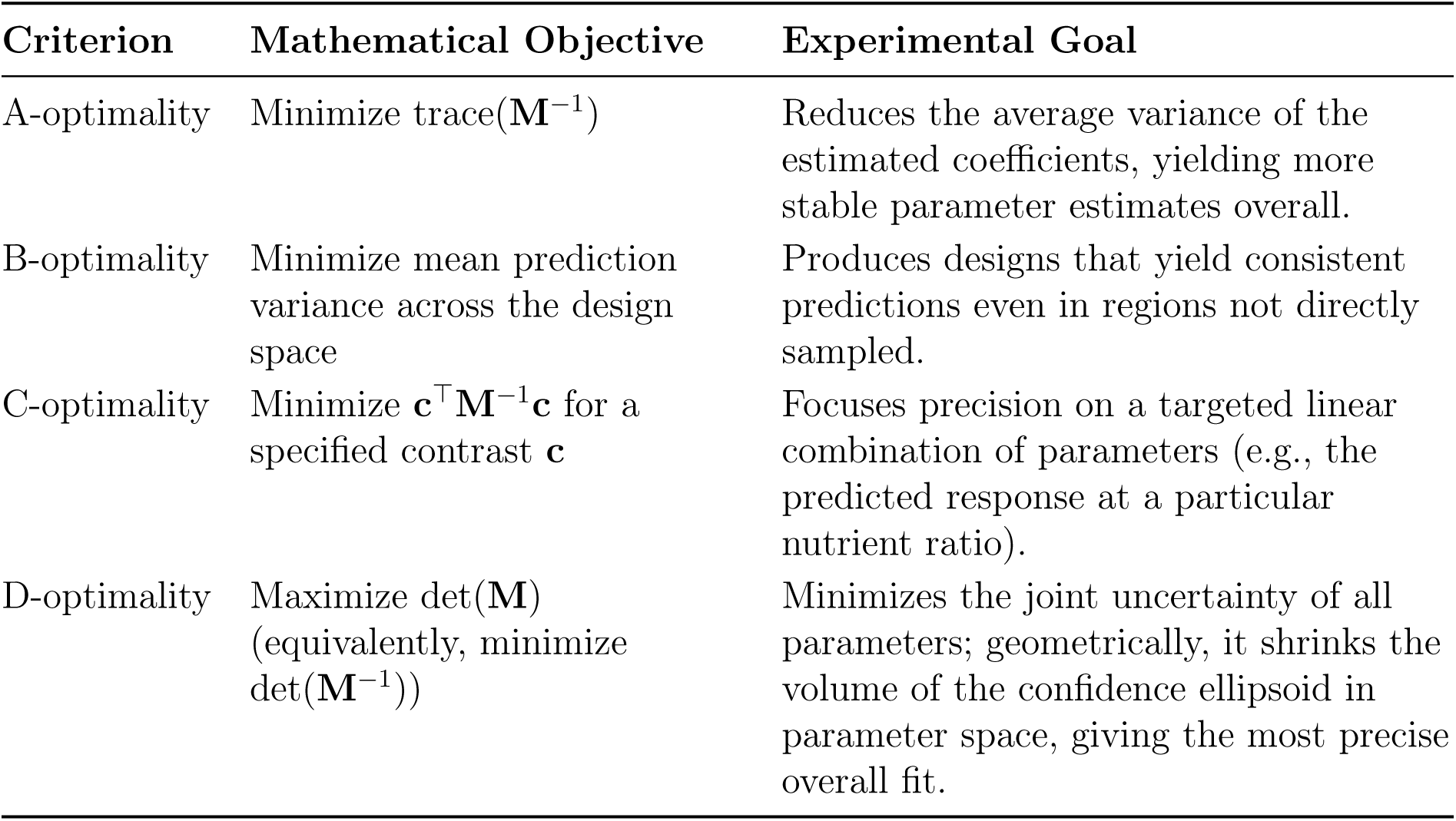
Summary of common optimality criteria used in mixture experiments (other criteria such as E, G, or I-optimality may also be appropriate depending on the objective; [69, 70]). Here, **M** denotes the Fisher information matrix of the fitted model over the chosen design; for linear models with homoskedastic errors, a standard choice is **M** = **X**^⊤^**X***/σ*^2^, where **X** is the design matrix and *σ*^2^ the error variance.

The implications of adopting the simplex framework span basic and applied domains. For instance, controlled nutrition studies are critical to ensuring that captive animals, such as laboratory rodents or managed insect colonies, receive balanced diets that support both health and well-being [32, 33, 34]. Animal production systems face increasing pressure to maximize growth and resilience while reducing feed costs [35]. Simplex-based mixture designs have been used to identify nutrient ratios that improve growth and feed efficiency in farmed species [36], as well as studies which utilize the GFN [37]. In experimental design, mixture models and response surface methodologies (RSM) provide a rigorous basis for disentangling competing hypotheses regarding nutrient interactions, maximizing power, and minimizing experimental cost [38, 39, 40, 41, 42, 43]. In ecological studies, simplex visualizations allow researchers to quantify how nutrient availability changes across trophic levels, revealing whether higher-order consumers are constrained by the nutrient spaces of their prey or transformed by synthetic processes [12, 10, 44]. Finally, in theoretical ecology, mixture triangle-based models can formalize foraging trajectories under alternative behavioral rules, enabling direct tests of optimal foraging theory [45, 46].

The simplex framework holds promise as a unifying scheme for experimental designs in nutritional ecology and applied animal sciences. By integrating visualization, mathematical modeling, and optimal experimentation, it provides a foundation for answering both fundamental and practical questions about how organisms acquire and regulate nutrition in complex choice and no-choice environments. Here, we expand on the framework by showing how concepts inherent to GFN (intake target, rules of compromise, etc.) manifest in this coordinate system, and we further develop some of its mathematical formalisms. We focus on statistical modeling of choice assays (where organisms can freely choose among a set of nutrient options over time) and no-choice experiments (where organisms are given a fixed diet; these are more generally known as mixture experiments), as well as the optimal design of experiments and how to perform multi-objective optimization of outcomes. We first compare and contrast the EMT and RMT coordinate systems (Section 2) and show how it can be used to design and analyze choice experiments (Section 3) and no-choice experiments (Section 4). We end with a discussion on the potential applications of the simplex in both a basic and applied setting (Section 5).

## 2. EMT and RMT Coordinate Systems

To first give an intuition as to how the EMT and RMT coordinate systems work, we show how to construct an EMT (Section 2.1), and how to make transformations between EMT and RMT to show their relations (Section 2.2). Next, we show how the RMT introduces visual bias that is not present in the EMT or simplex due to this transformation (Section 2.3).

### 2.1 Constructing a Simplex (Equilateral Mixture Triangle)

Mixture experiments often aim to understand how the proportions of different components combine to influence an outcome. In this framework we focus on the ratios of components rather than their absolute intake rates, which is appropriate when the question is about broad, lifetime dietary choices rather than the precise amounts required per unit time. While the amount of acquired nutrients can certainly be used to answer some ecological questions [3], to retain the visual intuition of the GFN in three dimensions, we must compress totals into ratios. To visualize these ratios, we begin with the simplest case: a mixture of two components.

For two components, every possible mixture can be represented along a single line (Figure 1A). One end of the line corresponds to 100% of the first component and none of the second, while the opposite end corresponds to 100% of the second and none of the first. Any point in between shows the ratio between them. For example, a mixture with equal parts of each component would lie halfway along this line. In this way, the space of all possible two-component mixtures can be thought of as one-dimensional, even though it represents combinations of two ingredients.

**Figure 1:**
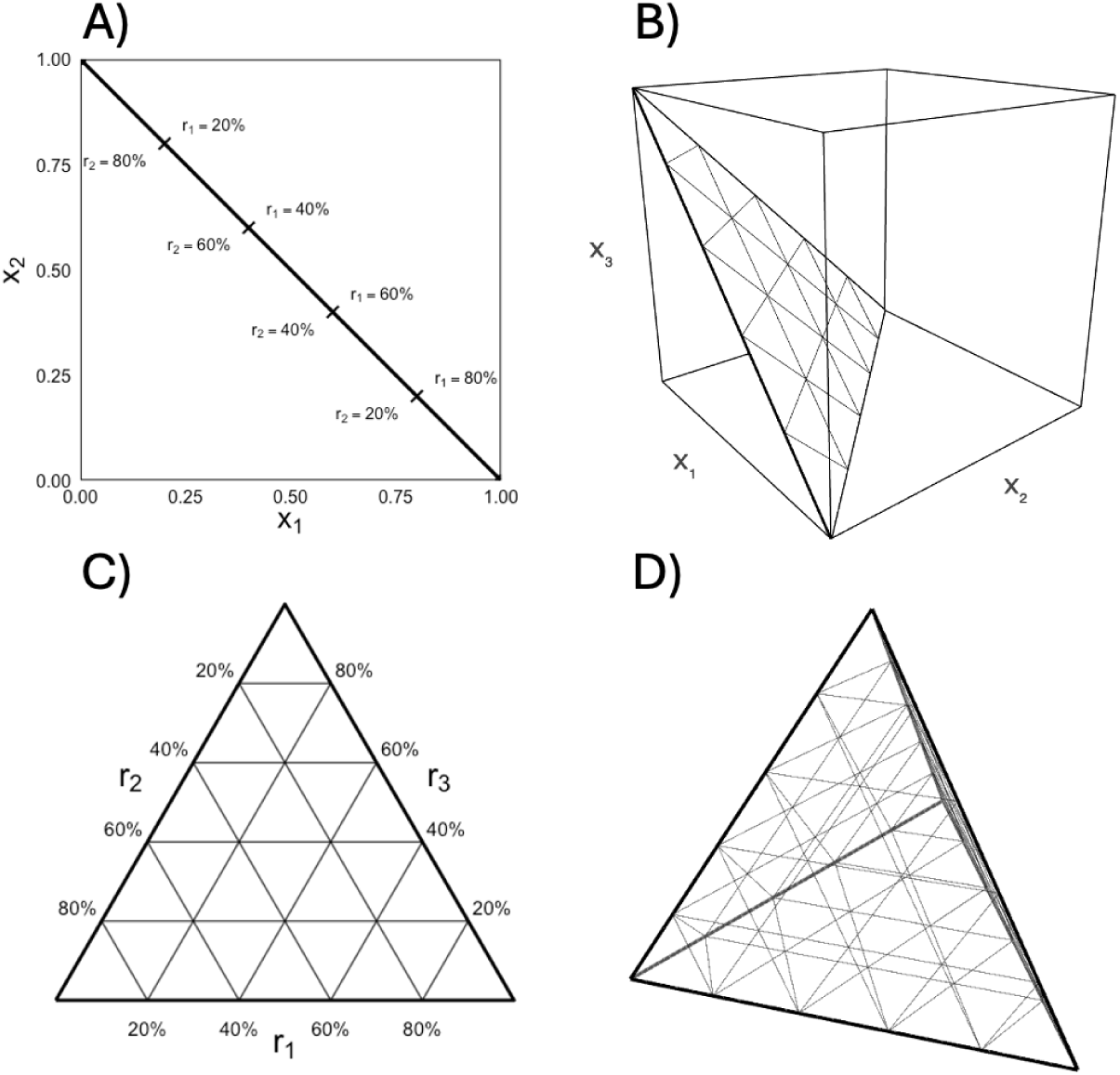
Illustration of the construction of the EMT. (A) One-dimensional simplex (binary mixture) shown as the diagonal from (0, 1) to (1, 0); tick marks give percentages of the two components *r*_1_ and *r*_2_. (B) The ternary slice inside the unit cube (white triangular face). (C) Equilateral ternary diagram of the same slice; external edge labels indicate *r*_1_ (bottom), *r*_2_ (left), and *r*_3_ (right). (D) Three-dimensional simplex (regular tetrahedron) for four-component mixtures. This can be thought of as a slice of a 4-dimensional hypercube.

Adding a third component expands on this idea. A three-component mixture must still sum to 100%, so the set of all possible combinations no longer fills a three-dimensional cube of possibilities—it lies within a two-dimensional slice of that cube. When drawn, this slice forms a triangle (Figure 1B). Each corner of the triangle represents a pure component (100% of one and 0% of the others), and every point inside the triangle corresponds to a specific combination of the three. The triangle is drawn with equal sides to make the distances between mixtures visually comparable, the EMT (Figure 1C). Moving toward one corner increases the proportion of that component while decreasing the others, allowing all three proportions to be viewed simultaneously in a single, balanced coordinate system.

This same logic generalizes beyond three components. A mixture with four components, for example, would occupy a three-dimensional region (Figure 1D), and one with five would occupy a four-dimensional region, and so on. Because mixtures are defined by proportions that sum to a constant, the space of all possible mixtures always has one fewer dimension than the number of components. The EMT is therefore a special case of a more general concept: the family of these shapes is called the simplex.

Higher-dimensional studies may be useful in some contexts. For example, since some species of fruit fly consume fermented fruit [47], ethanol may represent a fourth nutritional dimension, since it can also serve as another source of calories. Such studies would be trivial extensions of the simplex, and though is technically possible to perform using the RMT [16], the visualization is less interpretable in this latter case (and it is not clear how the RMT would extend past four dimensions).

A more formal definition of the coordinate system, given in the context of macronutrients, is provided in Supplemental Section S1, along with suggestions on how to model the nutritional state of an organism over time and how to model a series of foraging decisions. Some further applications of this modeling approach are given in Section 3.

### 2.2 Transformations Between Coordinate Systems

The EMT is not the only way to represent ratios for a three-component mixture. The RMT retains the same compositional information as the equilateral version, but it is plotted using a coordinate system that aligns with the standard horizontal and vertical axes. To give a better intuition as to how each coordinate system can be read and to better understand their visual differences, we show how one would transform an EMT to form an RMT.

In the equilateral representation, the three mixture components are arranged at the corners of an equilateral triangle, so that increasing the proportion of one component corresponds to movement directly toward its vertex. The right-angled version reorients this same mixture space. Instead of positioning components at equal angles, two components are placed along the horizontal and vertical axes, and the third component is defined implicitly, as the distance from the origin. That is, the closer a point is to the origin, the higher the ratio corresponding to the third component. A mixture that was originally represented inside the equilateral triangle is therefore “stretched” and “tilted” so that it fits into a right-angled coordinate frame. In this new view, mixtures are still defined by the same proportional constraints (the three parts must sum up to 100%), but now the contribution of each component can be read directly from the orthogonal axes.

To visualize the relationship between the two coordinate systems, we can imagine gradually deforming the equilateral triangle until one of its sides becomes horizontal and another becomes vertical (Figure 2). The process preserves the proportional structure of the mixture but redefines the visual geometry. Points, grid lines, and labels that were evenly spaced in the equilateral system are transformed into positions appropriate for the right-angled one (an affine transformation). Further, after this transformation, the orientation of the y-axis is flipped, which allows for a Cartesian interpretation of the RMT.

**Figure 2:**
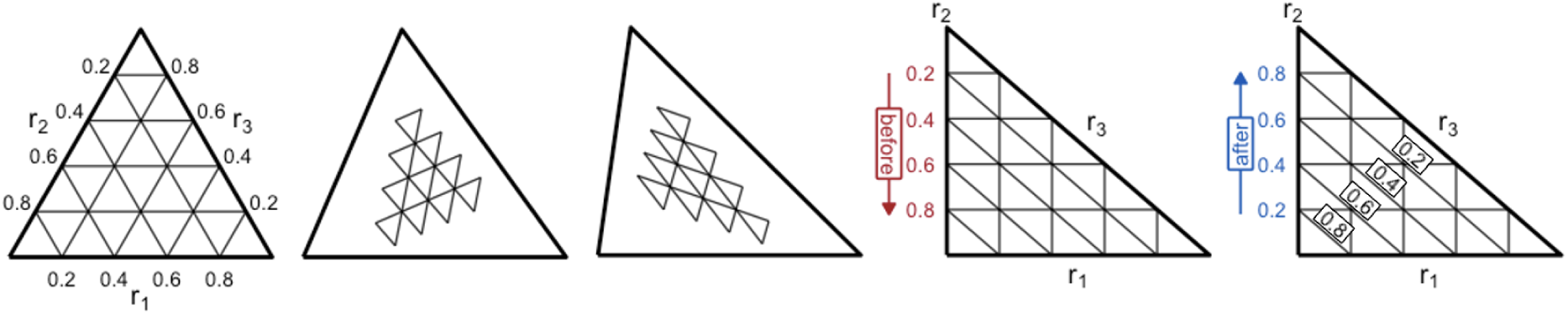
Transformation from an EMT to an RMT. Each panel shows a successive stage in the geometric transformation between two coordinate systems used to represent three-component mixtures. In the EMT (leftmost panels), the three components are positioned at the corners of an equilateral triangle, ensuring all are treated symmetrically and that changes in any component correspond to equal visual distances. As the transformation proceeds, the triangle is gradually stretched and reoriented so that two components align with the horizontal and vertical axes, producing the right-angled form (rightmost panels). At the end, the direction of the y-axis is reversed. This has the effect of converting the *r*_3_ axis into an implicit one, where ticks radiating from the origin to the hypotenuse now correspond to *r*_3_ and are ordered from high to low.

### 2.3 Visual Bias Introduced by the Right-Angled Mixture Triangle

Although the RMT conveniently displays mixture data using standard Cartesian axes, it can also introduce distortions that misrepresent the true relationships among samples.

As noted by [16], mixture plots are often used not only to display mean compositions but also to assess the scatter of replicate data within samples. For example, researchers have measured the variance in the macronutrient composition of milk in wild primates by plotting observations on a mixture triangle [19]. Thus when the coordinate system itself distorts distances or angles, this can create misleading impressions of variability or overlap between groups.

To illustrate this effect, we compared the same three sets of simulated data displayed in both the EMT and the RMT (Figure 3). Each group consists of an elliptical cloud of points representing replicate samples, with identical variation along their major and minor axes. In the EMT representation, these ellipses appear equal in size and shape, confirming that the amount of variation within each group is the same (Figure 3A). However, when the same data are projected into the RMT, the transformation preserves distances along the horizontal and vertical axes but compresses distances along the implicit diagonal axis by a factor of 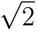 (Figure 3B; Supplemental Section S2). As a result, variation aligned with this diagonal appears artificially smaller or “squished,” even though the underlying variability is unchanged.

**Figure 3:**
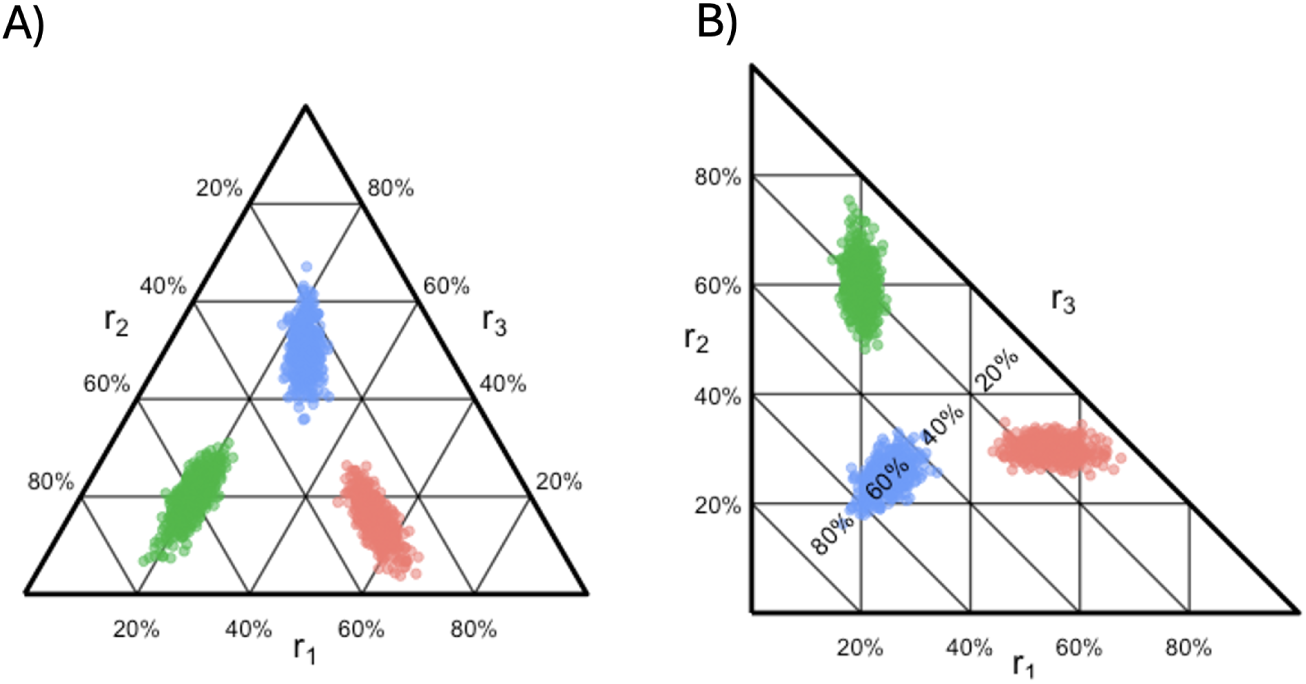
Comparison of variation among replicate samples in equilateral and right-angled mixture triangles. Each plot shows three groups of simulated mixture data, where each group has the same underlying amount of variation but with different orientations of its major and minor axes. A) In the EMT, these ellipses appear equal in size and shape, accurately reflecting the identical variability of the three groups. B) When the same data are displayed in the <u>R</u>MT, variation along the diagonal (the blue group) axis is compressed by a factor of 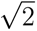, making one orientation appear artificially narrower. Diagonal ticks represent the levels of *r*_3_.

This geometric distortion undermines one of the key advantages of the GFN: its ability to visually convey balanced relationships among nutrients. While the RMT can be easier to construct and interpret using conventional plotting tools, its unequal scaling introduces a visual bias that can affect intuition about relative variation, overlap, and dispersion. The simplex, by contrast, preserves proportional relationships equally in all directions, ensuring that visual patterns correspond more faithfully to the true structure of the data.

## 3. Use of the Simplex for Choice Experiments

One of the most common uses for the GFN is in the evaluation of choice assays where animals are allowed to freely forage for two or more food items of differing nutritional content [48, 49, 50, 51, 52]. The principles used to construct GFN manifest differently when the total amount of nutrients is replaced by ratios in the coordinate space (see Section 2). We first discuss what the GFN looks like when plotted using a simplex (Section 3.1), introduce a novel hypothesis test to allow researchers to reject the null hypothesis of random choice (Section 3.2), and we reformulate the rules of compromise so that they are consistent with the simplex (Section 3.3). We end with a hypothetical case study showing how one would quantitatively measure and compare rules of compromise (Section 3.4).

### 3.1 Realizing the Geometric Framework for Nutrition within the Simplex

Many of the central ideas of the GFN can be represented naturally within the simplex. In the GFN, an organism’s nutritional state is typically described as its position within a state space defined by the amounts of multiple nutrients. Feeding behavior is then viewed as a trajectory toward an intake target (Figure 4): a point that represents the optimal balance of nutrients, independent of the total nutrient intake, that maximizes some fitness-relevant suite of traits (although other targets could exist as well, such as a growth target; [53]). In the wild, intake targets can often be defended due to the significant variation in nutrient compositions of food sources, for instance, in the prey of predatory seabirds [20]. Other species are highly specialized or have limited food choices, reducing the available nutrient sources that can be relied on for intake targets. Intake targets can be expected to assist in achieving a suite of compromising physiological, ecological, or ontogenic outcomes, such as minimizing development time [54] or maximize lifespan [55]. Within GFN, the intake target can be regarded as the total amount of nutrient consumed, which can be decomposed into a growth target (the fraction that is retained in new tissue or other beneficial stores), a nutrient target (the growth target plus the fraction of disassociated nutrient that is used for metabolism and other functions), and the remaining fraction of disassociated nutrient that is not used and is excreted or otherwise lost. Because absorption and utilization efficiencies can differ among nutrients, the macronutrient ratios associated with the growth and nutrient targets need not match those of the intake target itself, and this decomposition provides additional biological insight into how observed intake patterns relate to underlying physiological processes [10]. Both growth and nutrient targets also appear as points on the simplex.

**Figure 4:**
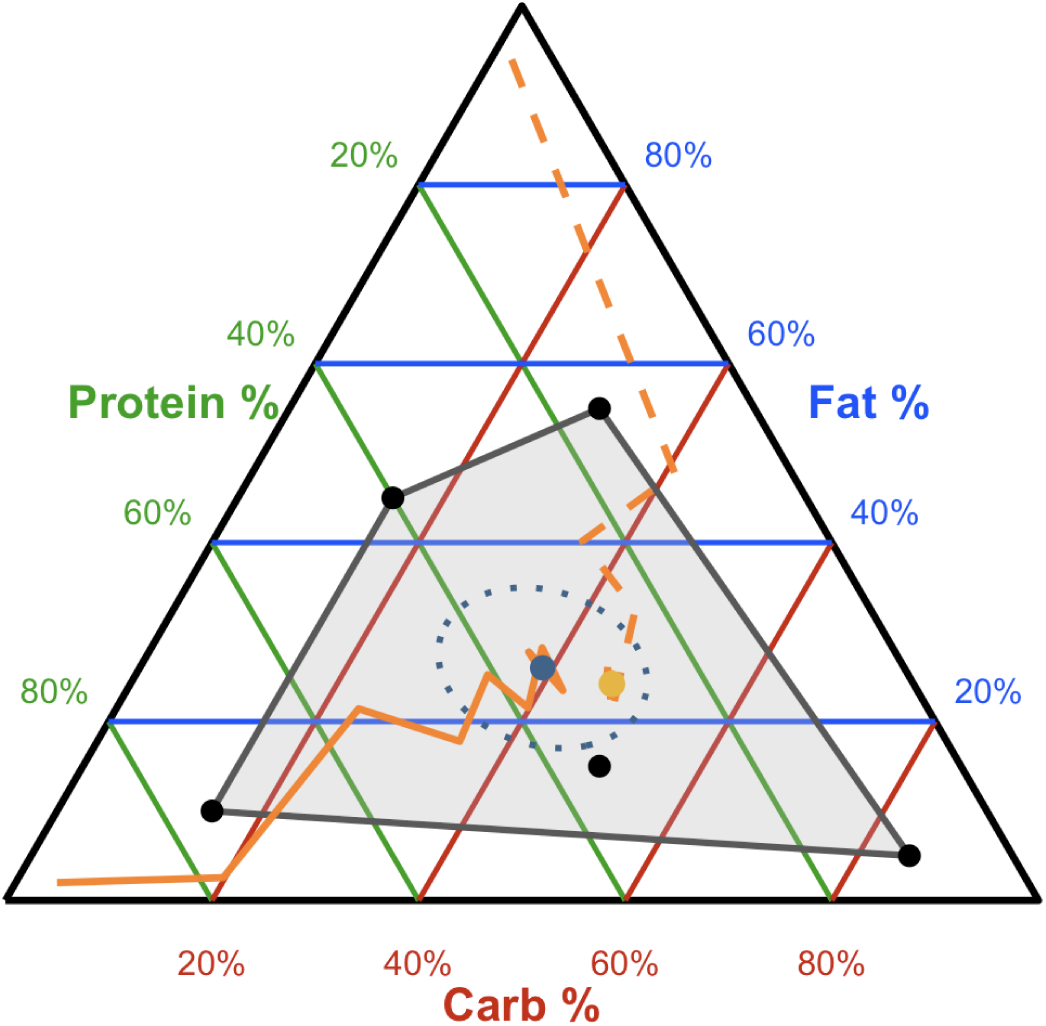
Key concepts of the GFN represented in the simplex. Black points mark the nutrient compositions of available foods, while the grey polygon connecting them (the convex hull) defines the set of all attainable mixtures, known as the nutrient space polygon. The blue point denotes the mean nutrient composition (center of mass) of all available foods, and the gold point indicates an intake target representing the ideal nutrient ratio for the organism. Orange trajectories illustrate two possible feeding paths through the space as an organism consumes different foods and gradually adjusts its nutrient balance. One trajectory represents random choice (solid orange line), while the other represents the defense of an intake target (dashed orange line). Once within the convex hull, the organism’s nutritional state cannot move outside this region through any combination of the available foods. The dotted blue line represents the 95% confidence interval for random choice. Food choices that end within this region likely cannot be distinguished from the random selection of food items (where every food item is equally likely to be selected). Here, the intake target is too close to the center of mass, so the defense of the target is indistinguishable from random choice.

Similarly, foods that were originally described as food rails (lines extending from the origin in the traditional GFN) are represented in the simplex as discrete food points. Each point represents the constant ratio of nutrients contained in a particular food. Feeding decisions can therefore be viewed as trajectories through the simplex. As an organism consumes foods in different ratios, its position in the triangle moves accordingly, updating its overall nutrient balance. The shape and direction of these trajectories reflect both the composition of the foods consumed and the behavioral rules governing food choice (for further discussion on how this can be done, see Sections S1.3 and S6). The set of all possible nutritional outcomes attainable from a given set of foods is defined by the smallest convex polygon that encloses all of the food points (on a traditional two-component GFN graph, this polygon corresponds to the full space between all available nutrient rails). This polygon is known as a convex hull, but it has been referred to as a nutrient space polygon in the GFN literature, where it is drawn on top of RMTs [56, 20, 57, 26]. While nutrient and growth targets will often lie close to the intake target in nutrient space, unequal absorption or post-ingestive processing of different nutrients can, in principle, shift these physiological targets to positions that fall outside the convex hull.

We demonstrate that the nutrient space polygon is indeed a convex hull (Supplemental Section S3). Thus, this boundary acts as an event horizon of sorts: once an organism’s nutritional state enters this region, it cannot move outside of it through any combination of those foods (unless a new food item is introduced or the nutritional content of an available item is transformed). This has strong experimental and theoretical implications which we explore in Supplemental Sections S1, S4, and S6.

### 3.2 Statistical Test for Random Choice

To evaluate whether observed feeding trajectories differ from what might occur by chance, we constructed a statistical baseline corresponding to random food choice (the null hypothesis in this context, [58, 59, 52]). Such a test is necessary when there is a complex nutritional landscape available to the test organism (which may be needed to test hypotheses using the GFN, [60]), so the null expectation may not be obvious as it would be in a binary choice assay, where the null expectation is that each option has a 50% probability of occurring [54]. In this null model, all foods are assumed to be equally likely to be chosen, such that the expected nutrient composition of a meal is simply the average of all available foods. Repeated random draws from these foods, therefore, yield a distribution of expected nutrient ratios that is centered at the mean composition (the blue point in Figure 4).

To visualize the variability associated with this random process, we estimate the two-dimensional variance–covariance structure of the nutrient ratios in simplex coordinates (derived in Section S4). Each food contributes to this variability according to its position relative to the mean, and the spread of points reflects the diversity of nutrient compositions among the available foods. Assuming each draw is independent, the uncertainty in the average composition decreases with the number of individual choices, following standard sampling theory.

From this covariance structure, we compute a 95% confidence ellipse that encloses the region in which 95% of the random-choice means would be expected to fall. The ellipse is derived by scaling the eigenvectors of the covariance matrix by the critical value of the chi-square distribution for two degrees of freedom. Its orientation and shape reflect both the relative variance along the major and minor axes of nutrient variation and the correlation between them.

This dotted ellipse (Figure 4) provides a useful visual benchmark: if the observed or simulated intake target lies well within the ellipse, the resulting nutrient balance could plausibly arise from random food choice. Conversely, if the intake target or feeding trajectories fall outside this region, the pattern likely reflects directed regulation rather than stochastic sampling. Experiments, therefore, should be designed such that the expected intake target does not overlap this ellipse. This may require changing the geometry of the convex hull by providing different combinations of foods, or it may mean that animals should be allowed to make more foraging decisions so that the axes of the ellipse can shrink.

Although the geometric approach described above captures the basic layout of the available foods, it can sometimes produce an ellipse that extends into parts of nutrient space the organism could never actually reach. In other words, the mathematical shape may spill outside the true boundary formed by the foods on offer. A more realistic way to estimate where an organism might end up is to use a simple simulation. The idea is to repeatedly mimic what would happen if the organism chose foods at random, doing so the same number of times it actually foraged (or the number of times the experimenter allows it to forage). Each simulated sequence of choices produces a final nutrient state. By repeating this process many times, we obtain a cloud of possible outcomes. We can then draw the smallest possible polygon that contains 95% of these points, giving us an empirical 95% region that reflects realistic boundaries based solely on the foods the organism could truly reach.

### 3.3 Rules of Compromise in Simplex

In the GFN, the rules of compromise describe how an animal adjusts its feeding behavior when confronted with nutritionally imbalanced or suboptimal food choices. These rules are typically formulated in terms of absolute nutrient amounts (nutrient arrays, which are the collection of average food intake of animals in each of the nutritional rails; [61]), for instance, how far an animal deviates from its intake target along each nutrient axis when only certain foods are available [62, 60, 8]. However, when we move from an amount-based coordinate system to a ratio-based one, such as the EMT or the RMT, these traditional compromise rules cannot be directly mapped. This is because ratio spaces preserve only relative proportions among nutrients, not their total quantities, so the geometries of nutrient arrays are effectively erased. Despite this, one can still visualize how animals contend with suboptimal mixtures by examining trajectories through nutrient space rather than absolute amounts.

In the simplex, each point represents a nutrient ratio rather than a total intake. Thus, the process of compromise can be visualized as a path through this ratio space that reflects how the organism’s nutrient balance changes with successive feeding decisions. When multiple food options are available, one of the primary rules of compromise, closest distance optimization, can be realized as a direct trajectory between the organism’s current nutritional state and its intake target. This path represents the shortest route through nutrient space to the ideal nutrient ratio, but in practice it is often choppy, as the animal must move incrementally between the nutrient compositions of the available foods (Figure 5A; see Section S1.4 for a conjecture regarding how this trajectory may be derived).

**Figure 5:**
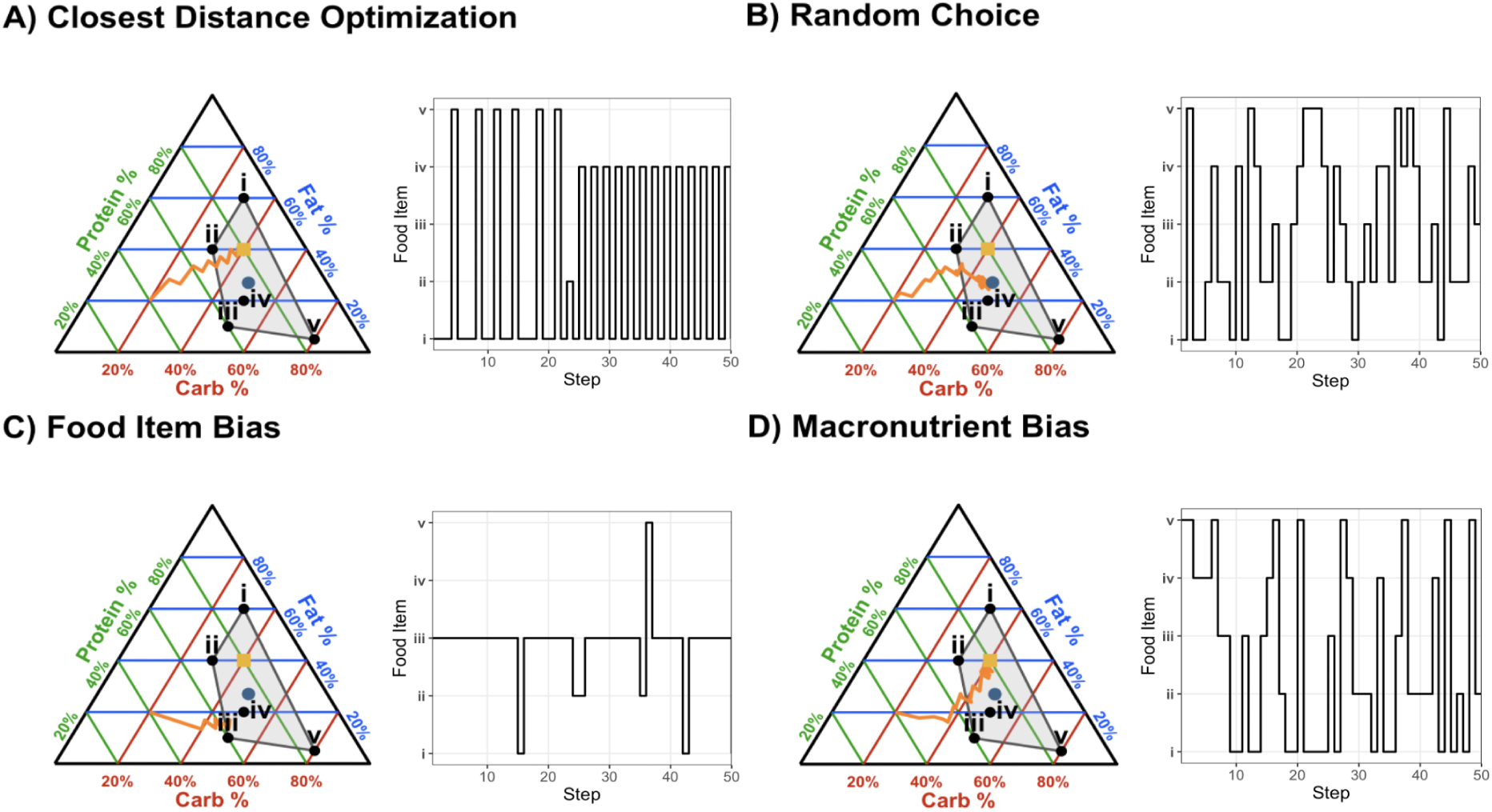
Compromise trajectories in the Simplex. The left side of each panel shows a feeding trajectory (orange path) from an initial nutrient ratio toward either an intake target (gold square) or a limiting point determined by the decision rule in the simplex. The black dots (i–v) are available foods; the gray polygon is the convex hull (the possible nutrient space); the blue dot is the center of mass (mean composition) of all foods. Axes are colored by nutrient: Protein (green, left), Fat (blue, right), Carbohydrate (red, bottom). The right side of each panel shows the time series of chosen foods. A) Represents food choice based on closest-distance optimization. At each step, the food that yields the smallest next-step distance to the intake target is chosen, producing a piecewise “choppy” approximation to the direct path. B) represents random choice, or uniform selection among foods, causes convergence to the foods’ center of mass. C) represents a food-item bias: preference for a specific item (here, food *iii* ) drives convergence to that item. D) gives a macronutrient bias, where the organism will first converge on the optimal ratio of one macronutrient (here, carbohydrates) before attaining the optimal ratios of the other two macronutrients.

If the organism exhibits no preference among the available foods, each food item being equally likely to be consumed, then the expected trajectory will converge upon the center of mass of the convex hull formed by the food points (Figure 5B). This outcome reflects the mean composition of the available options and corresponds to a purely random choice strategy. Conversely, if one food item is preferred over another (for example, if one food source lies closer to a central-place forager’s nest and therefore minimizes travel cost) then optimal foraging theory predicts that the organism’s trajectory will converge toward that particular food (Figure 5C; [63]). Finally, if the organism immediately requires one nutrient than the others, then its trajectory is expected to initially move along the axis representing that nutrient before gradually converging toward the intake target. In Figure 5D, for instance, the organism prioritizes carbohydrate balance before adjusting the ratios of protein and fat. A general framework for modeling each of these foraging decisions is given in Section S1.3.

Generally, it can be difficult to assess the rule of compromise from the time series of decisions alone (Figure 5, right side of panels). However, knowing the time order of observations can be useful. For instance, if an organism is behaving optimally, then researchers may know that she reached her intake target because of a consistent pattern in food choices (Figure 5A, second panel). This could be used as a decision rule to terminate an experiment early, saving time and resources.

### 3.4 Case Study: Using the Simplex to Infer Rules of Compromise

To illustrate how the simplex framework can be used to infer behavioral rules of compromise, we consider a hypothetical choice experiment designed to determine which decision rule an organism employs to defend its nutritional intake target when no single food item provides the ideal macronutrient ratio. In this experiment, the organism is presented with five food items that have distinct proportions of carbohydrate, protein, and fat. Each of these foods has been confirmed as safe for consumption and falls within the nutrient space polygon (the convex hull) defined by the mixture of available foods.

Prior to the main trial, the organism is offered a single priming food that lies outside of this convex hull, containing a composition far from the target balance of nutrients. This priming phase ensures that the organism begins the trial in a nutritionally imbalanced state, making the subsequent trajectory through nutrient space clearly observable (Figure 6A). After this initial feeding, the organism is placed in an arena where all five experimental foods are available simultaneously.

**Figure 6:**
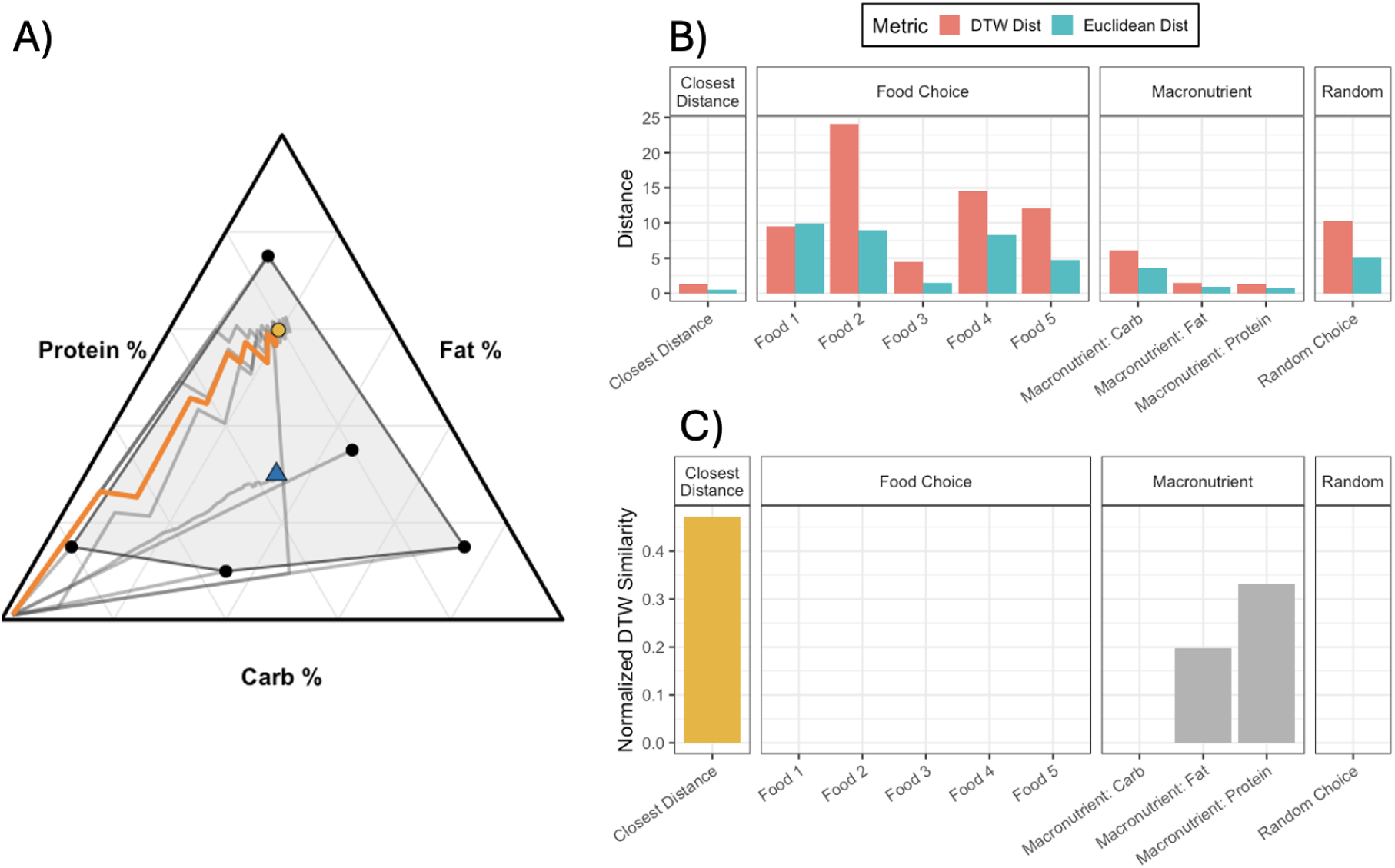
Comparing real nutritional trajectories to theoretical trajectories of various rules of compromise. A) The simplex showing the five available foods (black circles), their convex hull (gray region), the center of mass (blue triangle), the theoretical trajectories from various rules of compromise (gray lines), the intake target (gold circle), and the observed trajectory of the organism through nutrient space (orange line). B) Grouped bar plot comparing raw distance measures (Euclidean and dynamic time warping, DTW) across each theoretical rule of compromise, with categories corresponding to random choice, single-food preference, macronutrient prioritization, and closest-distance optimization. C) Bar plot showing the normalized similarity scores derived from dynamic time warping between the observed trajectory and the same rules of compromise. The gold bar shows the category with the tightest correspondence to the true trajectory. Together, these panels demonstrate how the simplex framework can be used to infer the decision rule that best describes an organism’s foraging behavior.

The organism is then allowed to forage freely until it either ceases feeding or begins repeating a predictable sequence of food choices. Either outcome may indicate that the organism has reached its intake target or achieved homeostasis in nutrient balance. Throughout the trial, each feeding bout is recorded and used to reconstruct the organism’s nutritional trajectory through time. These cumulative nutrient states are mapped onto the simplex as a continuous path, shown as the orange line in Figure 6. This trajectory represents the animal’s observed movement through nutrient space, beginning from its primed state and terminating near its intake target.

To interpret this observed trajectory, it is compared against several theoretical trajectories that represent alternative decision strategies, or rules of compromise (Section 3.3). The mathematical formulation of these trajectories, their underlying decision rules, and the methods by which we compare trajectories is provided in Supplemental Section S6. Each theoretical path can be treated as a reference model, and the observed trajectory can be quantitatively compared to each by measuring the distance between their paths on the simplex (Figure 6B). These distances are then transformed into normalized similarity scores using an exponential decay function (Figure 6C). This scoring system translates raw distances into interpretable measures of correspondence between the observed behavior and each theoretical rule of compromise. In this hypothetical example, the highest similarity score corresponds to the “closest distance optimization” rule, suggesting that the organism’s foraging decisions most closely follow a deterministic strategy that minimizes the distance to the intake target at each step.

Beyond this illustrative case, the experimental framework can be extended to explore the consistency and generality of decision rules. For instance, would the same organism exhibit the same strategy if presented with a different set of food items that define a new nutrient space? Would it still follow the shortest path toward the target, or would its choices depend on the configuration of available foods? These extensions allow researchers to test whether rules of compromise are objective, intrinsic decision-making heuristics, or instead emergent properties determined by the nutritional environment or the organism’s internal states. The simplex framework thus provides a unified geometric and probabilistic approach to studying nutritional decision-making across contexts.

## 4. Use of the Simplex for No-Choice Experiments

In no-choice experiments, individuals or groups are restricted to a single diet or resource mixture for a single run, allowing researchers to quantify how performance or physiological traits respond to variation in composition alone. These experiments are equivalent to mixture experiments performed in various engineering fields [64]. Thus, these experiments can be performed within the same theoretical framework as choice experiments.

In the following sections, we describe how no-choice assays can be optimized and interpreted within this framework. We first discuss principles of optimal experimental design that ensure efficient coverage of the nutrient space (Section 4.1). We then address standard statistical modeling in simplex, emphasizing mixture-model formulations that respect the compositional constraint (Section 4.2). Finally, we consider multi-objective optimization, where competing biological or experimental objectives (such as maximizing performance while minimizing cost) are jointly analyzed to identify trade-offs and Pareto-optimal nutrient ratios (Section 4.3). We also provide a simulated case study on what we suggest is an appropriate workflow for these types of experiments (Section 4.4) and summarize recommendations for both choice and no choice experiments in Table 3.

**Table 3:**
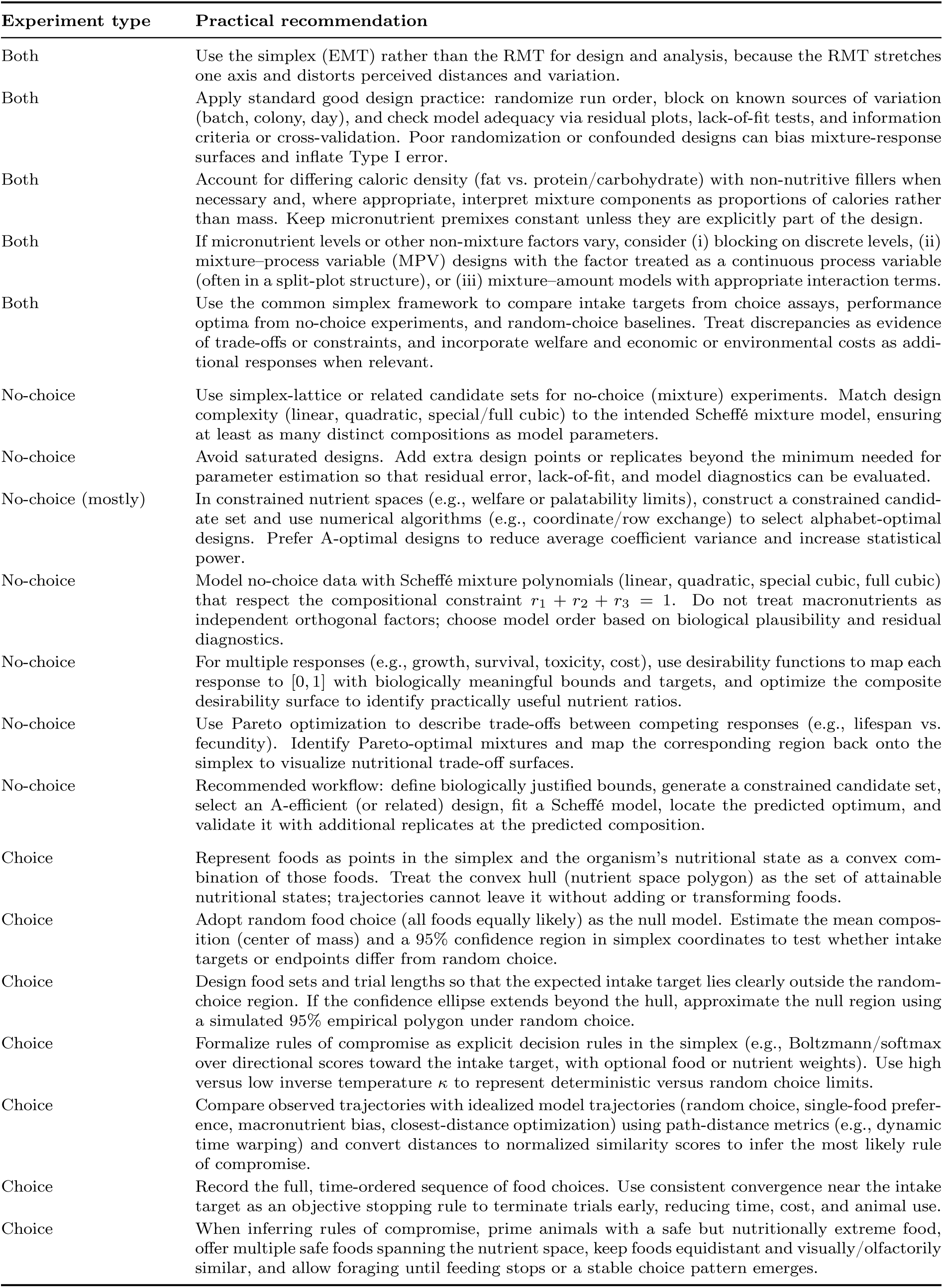
Practical experimental recommendations for choice and no-choice (mixture) experiments in the simplex (EMT) framework.

| Experiment type | Practical recommendation |
| --- | --- |
| Both | Use the simplex (EMT) rather than the RMT for design and analysis, because the RMT stretches one axis and distorts perceived distances and variation. |
| Both | Apply standard good design practice: randomize run order, block on known sources of variation (batch, colony, day), and check model adequacy via residual plots, lack-of-fit tests, and information criteria or cross-validation. Poor randomization or confounded designs can bias mixture-response surfaces and inflate Type I error. |
| Both | Account for differing caloric density (fat vs. protein/carbohydrate) with non-nutritive fillers when necessary and, where appropriate, interpret mixture components as proportions of calories rather than mass. Keep micronutrient premixes constant unless they are explicitly part of the design. |
| Both | If micronutrient levels or other non-mixture factors vary, consider (i) blocking on discrete levels, (ii) mixture-process variable (MPV) designs with the factor treated as a continuous process variable (often in a split-plot structure), or (iii) mixture-amount models with appropriate interaction terms. |
| Both | Use the common simplex framework to compare intake targets from choice assays, performance optima from no-choice experiments, and random-choice baselines. Treat discrepancies as evidence of trade-offs or constraints, and incorporate welfare and economic or environmental costs as additional responses when relevant. |
| No-choice | Use simplex-lattice or related candidate sets for no-choice (mixture) experiments. Match design complexity (linear, quadratic, special/full cubic) to the intended Scheffé mixture model, ensuring at least as many distinct compositions as model parameters. |
| No-choice | Avoid saturated designs. Add extra design points or replicates beyond the minimum needed for parameter estimation so that residual error, lack-of-fit, and model diagnostics can be evaluated. |
| No-choice (mostly) | In constrained nutrient spaces (e.g., welfare or palatability limits), construct a constrained candidate set and use numerical algorithms (e.g., coordinate/row exchange) to select alphabet-optimal designs. Prefer A-optimal designs to reduce average coefficient variance and increase statistical power. |
| No-choice | Model no-choice data with Scheffé mixture polynomials (linear, quadratic, special cubic, full cubic) that respect the compositional constraint $r_1 + r_2 + r_3 = 1$. Do not treat macronutrients as independent orthogonal factors; choose model order based on biological plausibility and residual diagnostics. |
| No-choice | For multiple responses (e.g., growth, survival, toxicity, cost), use desirability functions to map each response to $[0, 1]$ with biologically meaningful bounds and targets, and optimize the composite desirability surface to identify practically useful nutrient ratios. |
| No-choice | Use Pareto optimization to describe trade-offs between competing responses (e.g., lifespan vs. fecundity). Identify Pareto-optimal mixtures and map the corresponding region back onto the simplex to visualize nutritional trade-off surfaces. |
| No-choice | Recommended workflow: define biologically justified bounds, generate a constrained candidate set, select an A-efficient (or related) design, fit a Scheffé model, locate the predicted optimum, and validate it with additional replicates at the predicted composition. |
| Choice | Represent foods as points in the simplex and the organism’s nutritional state as a convex combination of those foods. Treat the convex hull (nutrient space polygon) as the set of attainable nutritional states; trajectories cannot leave it without adding or transforming foods. |
| Choice | Adopt random food choice (all foods equally likely) as the null model. Estimate the mean composition (center of mass) and a 95% confidence region in simplex coordinates to test whether intake targets or endpoints differ from random choice. |
| Choice | Design food sets and trial lengths so that the expected intake target lies clearly outside the random-choice region. If the confidence ellipse extends beyond the hull, approximate the null region using a simulated 95% empirical polygon under random choice. |
| Choice | Formalize rules of compromise as explicit decision rules in the simplex (e.g., Boltzmann/softmax over directional scores toward the intake target, with optional food or nutrient weights). Use high versus low inverse temperature $\kappa$ to represent deterministic versus random choice limits. |
| Choice | Compare observed trajectories with idealized model trajectories (random choice, single-food preference, macronutrient bias, closest-distance optimization) using path-distance metrics (e.g., dynamic time warping) and convert distances to normalized similarity scores to infer the most likely rule of compromise. |
| Choice | Record the full, time-ordered sequence of food choices. Use consistent convergence near the intake target as an objective stopping rule to terminate trials early, reducing time, cost, and animal use. |
| Choice | When inferring rules of compromise, prime animals with a safe but nutritionally extreme food, offer multiple safe foods spanning the nutrient space, keep foods equidistant and visually/olfactorily similar, and allow foraging until feeding stops or a stable choice pattern emerges. |

### 4.1 Applying Mixture Experiment Principles to Nutritional Experiments

#### 4.1.1 Design and Construction of Mixture Experiments

Designing no-choice experiments in the context of mixture systems requires balancing experimental feasibility with the need to characterize the response surface across the simplex accurately. Because all component proportions must satisfy the compositional constraint *r*_1_ + *r*_2_ + *r*_3_ = 1, experimental points cannot be chosen arbitrarily within a Cartesian space; they must instead be selected from within the simplex. The goal of an optimal design is to select mixtures that provide maximal information about the response surface given practical constraints on sample size and the allowable range of compositions. Well-chosen experimental designs yield smooth, interpretable response surfaces that can reveal nutrient interactions such as trade-offs, synergies, or saturation effects.

A common starting point for constructing mixture designs is the simplex–lattice (Figure 7A) or simplex–centroid design. In a simplex–lattice design of degree (the number of components) *q* for *k* + 1 levels, each component proportion can take on values in the dis-crete set {0, 1*/k,* 2*/k, … ,* 1} , subject to the constraint that all proportions sum to one. The design is commonly denoted as *{q, k}*, producing 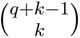 possible mixtures (which gives the sample size *n* for single-replicate designs). These points form a uniform grid of candidate compositions evenly distributed throughout the simplex, ensuring balanced coverage of the nutrient space. The simplex–centroid design, in contrast, focuses on all possible averages of the component vertices, producing a smaller but symmetric set of mixtures. Lattice designs are widely used because they generate aesthetically smooth and statistically stable response surfaces and because they generalize naturally to more than three components (*q >* 3), scaling well with increasing experimental complexity.

**Figure 7:**
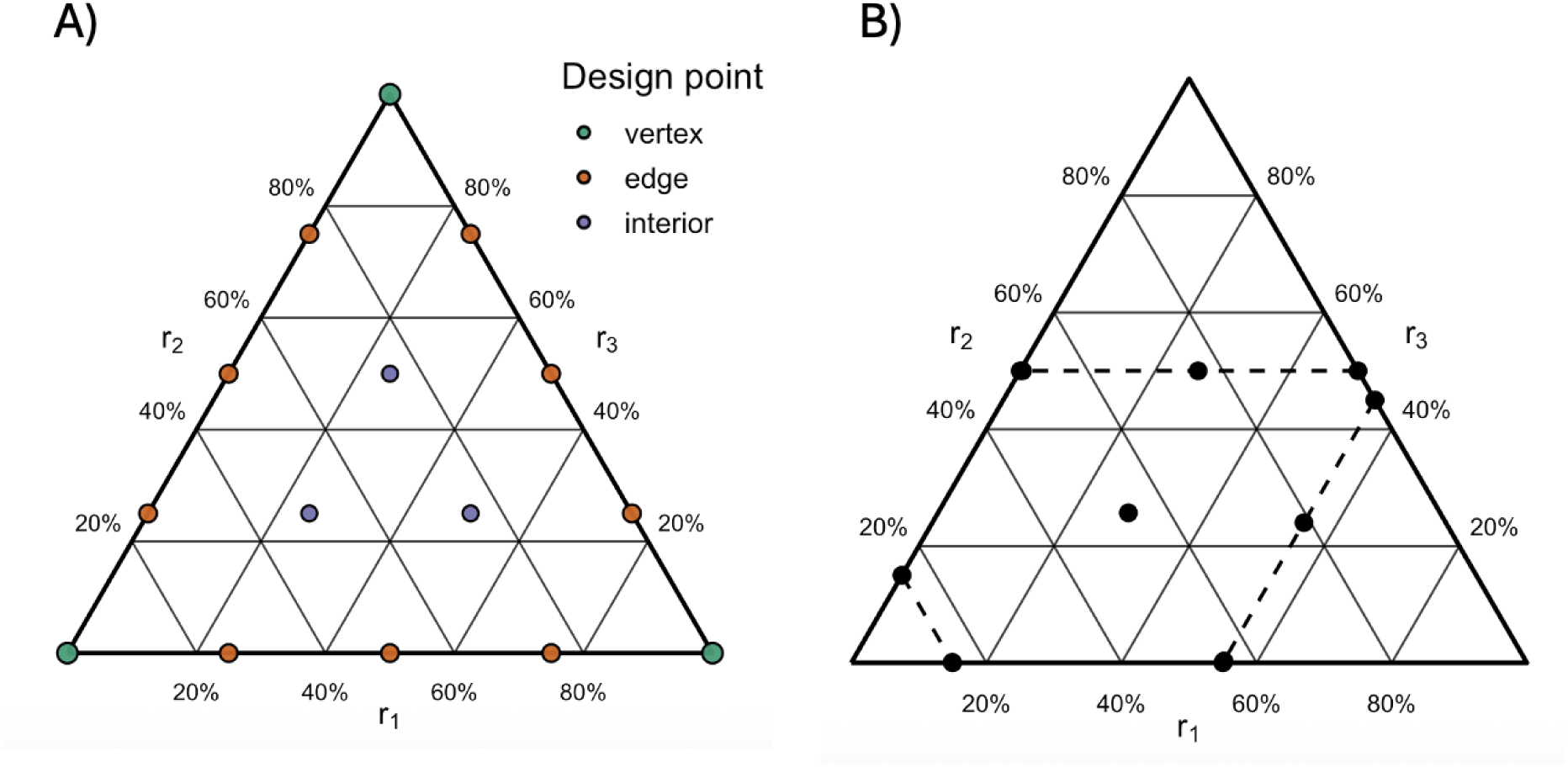
Examples of the design of mixture experiments. A) An example of a 3, 4 lattice design on a simplex, with the different types of design points marked in different colors. Extreme design points where only one macronutrient is present are known as a vertex. Points where two macronutrients are present are called edges. Finally, points where all three macronutrients are present are referred to as interior points. B) An example of a constrained mixture design, where design points were chosen to maximize D-optimality.

Additionally, the chosen statistical model data will dictate the smallest possible sample size. See Section 4.2.2 for the minimal sample sizes of commonly used mixture regression models.

#### 4.1.2 Optimality and Statistical Efficiency

While canonical lattice designs provide systematic coverage, they are not always realistic when resources are limited or when biological or logistical constraints restrict the feasible range of nutrient mixtures. For instance, in mice, extremely low protein diets can cause weight loss, rectal prolapse, or failure to thrive [65], consistent with studies of crickets where the recommended minimum protein content is 30% [66]. In such cases, optimal design theory provides a principled means of selecting the most informative subset of candidate points [67, 68]. Optimal design theory gives the set of what is known as alphabet-optimal designs, where each letter (i.e., A, B, or C) is the signifier for a different optimality criterion. These optimality criteria quantify how efficiently a design estimates model parameters or predicts responses (Figure 7B). These criteria are computed from the experimental design matrix **X** (see supplemental section S5 for an explanation of this matrix) and the Fisher information matrix **M**, which summarizes how much information each design point contributes about the model parameters. Table 2 summarizes some of the most common forms of optimality used in mixture experiments.

From an intuitive standpoint, minimizing variance (whether for individual coefficients, combinations of coefficients, or their joint confidence region) is desirable because variance represents uncertainty. Smaller variances indicate that the fitted model provides sharper, more reliable estimates of how the response surface behaves, and that predictions based on these estimates will be less sensitive to random experimental noise. For biological systems, this translates to greater confidence in inferred nutrient optima and in the shapes of trade-offs that define organismal performance. Maximizing A-efficiency (the degree to which A-optimality has been achieved) in particular has been shown to also maximize statistical power, which reduces the need for larger sample sizes [40].

When the design region is constrained (such as by nutritional irrelevance to the conditions of interest or limits on animal tolerance), these optimality criteria can be evaluated over the feasible subset of the simplex. Numerical optimization algorithms (such as coordinate exchange algorithms) can then be used to identify the subset of mixtures that provides the highest statistical efficiency under these constraints. The resulting designs need not be evenly spaced: they strategically concentrate sampling effort in regions that maximize the quality of the fitted statistical model. Such designs ensure that no-choice assays are both efficient and interpretable, producing response surfaces that faithfully represent the underlying nutrient-performance relationships.

#### 4.1.3 Practical Experimental Considerations

Both choice and no-choice experiments must also adhere to the general principles of sound experimental design to ensure that statistical inferences drawn from mixture studies are valid [71, 27]. Randomization minimizes bias due to uncontrolled factors, while blocking can account for known sources of variation such as batch, colony, or day effects. Model adequacy should always be assessed through residual diagnostics, lack-of-fit tests, and validation on replicate points. Additionally, competing models should be compared using information criteria or cross-validation to prevent overfitting. Attention to these fundamentals of the design is critical, as poorly randomized designs or poorly constructed designs (those that may accidentally conflate effects) can distort parameter estimates [72], inflate Type I error rates [73], and generate false nutrient–response relationships that could misrepresent underlying biological processes [74].

Another important consideration in experimental design is the differing caloric densities of macronutrients. Fat provides approximately 9 kcal g^−1^, whereas protein and carbohydrate each provide about 4 kcal g^−1^ [75]. To ensure that the effects of nutrient composition are not confounded by differences in caloric density, it may be necessary to incorporate a non-nutritive filler (e.g., indigestible cellulose; [76]). In such cases, the mixture components should be interpreted not as ratios of total mass, but rather as ratios of total caloric content.

For mixture experiments, essential nutrient concentrations should be held equal across all design points to prevent confounding, unless variation in these micronutrients is an explicit experimental factor. When essential nutrient variation is of interest, several design strategies are available. (i) Blocking: treat discrete levels of the essential nutrient (or preparation batch) as blocks and randomize mixture runs within blocks [70]. (ii) Mixture–process variable (MPV) designs: handle the essential nutrient as a non-mixture (continuous) process variable crossed with the mixture, using MPV models and ABC-optimal designs; this approach estimates mixture–process interactions and is often analyzed with split-plot structures when the process variable is harder to change [77, 78, 79]. (iii) Mixture–amount or covariate formulations: when total amount or a continuous supplement level covaries with the mixture, use mixture–amount models or include the factor as a covariate with appropriate interaction terms [77, 80]. In practice, MPV or split-plot MPV designs generally provide greater efficiency and interpretability than simple blocking when the micronutrient is continuous and expected to interact with composition.

### 4.2 Statistical Modeling of Mixture Experiments

#### 4.2.1 Introduction to Mixture Modeling

Although the RMT has been used to visualize macronutrient ratios as if they were orthogonal, the underlying proportions in any mixture system are not statistically independent. Because all components must sum to one, an increase in one proportion necessarily entails a decrease in at least one other. This constraint violates the assumption of orthogonality that underlies most factorial designs. Orthogonality refers to a condition in which explanatory variables are uncorrelated, ensuring that the estimated effect of one factor is independent of the others. In mixture experiments, this condition cannot be satisfied: macronutrient proportions are inherently collinear. However, it is a misconception that this dependence prevents the inclusion of all three nutrients as explanatory variables in a statistical model [25]. Rather, it requires that such variables be incorporated within a compositional or mixture model that explicitly accounts for their interdependence.

Mixture and compositional models were developed specifically to address this lack of orthogonality [70, 39]. These frameworks rely on transformed variables or constrained polynomial representations that ensure predictions remain valid within the simplex defined by *r*_1_+*r*_2_+*r*_3_ = 1. Among the most widely used are the Scheffé polynomials, which express the response as a function of component proportions and their products (e.g., *r*_1_*r*_2_, *r*_1_*r*_3_, *r*_2_*r*_3_), allowing for interaction and curvature effects while preserving the compositional constraint.

A response surface represents the prediction landscape of a model that has been fit to experimental data, typically using polynomial regression [39, 38]. In mixture experiments, it describes how a biological or performance variable (e.g., growth rate or reproductive success) changes as a function of nutrient proportions. Well-chosen experimental designs yield smooth, interpretable response surfaces that can reveal nutrient interactions such as trade-offs, complementarity, or limiting factors. In nutritional experiments, these surfaces can indicate whether an organism’s performance is optimized along a ridge (indicating substitutable nutrients) or peaked at a point (indicating synergistic effects).

#### 4.2.2 Scheffé Models of Increasing Order

Here, we present the most commonly used Scheffé models for mixture experiments in order of their complexity. We start with a linear model, then a quadratic model, and finally we present two cases of a cubic model. In these models, the coefficients *β_i_*represent the pure–component effects, meaning that *β_i_* corresponds to the expected response associated with the *i*th component in isolation (or its proportional contribution in the canonical mixture polynomial). The *δ* coefficients allow for synergistic or antagonistic blending effects.

For a mixture of *q* components, the simplest linear Scheffé model assumes purely proportional effects:

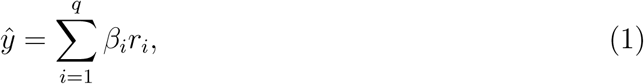

which describes a planar surface where each component contributes independently to the response. The minimal number of experimental points required to estimate this model is *n*_min_ = *q*. For three components, this corresponds to *n*_min_ = 3.

The quadratic Scheffé model introduces binary blending (pairwise interaction) terms, allowing for curvature and blending effects:

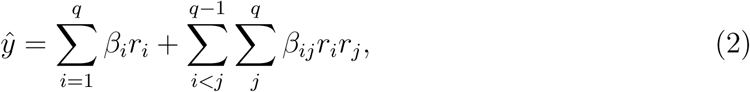

which, for three components, yields six coefficients and thus requires at least

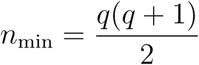

which equals 6 for a three-component system. Quadratic models can capture convex, concave, or saddle-shaped surfaces within the mixture triangle.

The special cubic Scheffé model extends this further by including selected ternary blending (third-order interaction) terms while maintaining model parsimony:

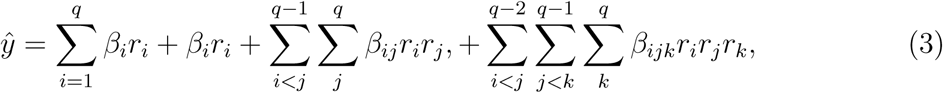

which allows nonlinear interactions between pairs of components but omits the full three-way term *r*_1_*r*_2_*r*_3_. The minimal number of experimental points required to estimate all parameters in a special cubic Scheffé model with *q* components is given by

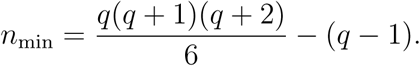

For a three-component system (*q* = 3), this yields *n*_min_ = 8 after applying the compositional constraint *r*_1_ + *r*_2_ + *r*_3_ = 1, which removes redundant parameters.

Finally, the full cubic Scheffé model includes all three-way interaction terms, providing the most flexible polynomial capable of describing highly nonlinear blending behavior:

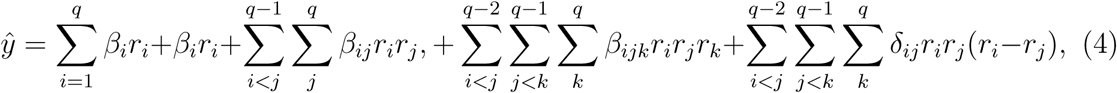

which requires

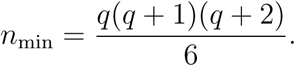

For three components, this corresponds to *n*_min_ = 10.

It is important to note that the sample sizes above correspond to saturated designs, meaning that the number of experimental points equals the number of model parameters. In such cases, residual error cannot be estimated (and thus, hypothesis testing), model adequacy checks, and lack-of-fit tests cannot be performed. For this reason, it is recommended that researchers include additional degrees of freedom by adding replicated or strategically distributed design points within the simplex to allow for estimation of experimental error and model validation.

#### 4.2.3 Interpretation in Nutritional Contexts

These polynomial models allow estimation of response surfaces that may be convex, concave, or saddle-shaped within the simplex. In the GFN, such surfaces may adopt a variety of topologies (flat, peaked, ridged, or curved), each corresponding to a different biological regime of nutrient interaction [6]. The accuracy of these surfaces depends critically on the number and spatial distribution of experimental design points within the mixture space. Poorly distributed or undersampled designs can bias parameter estimates and distort the inferred geometry of the nutritional response. Generally, polynomial regressions, as used in response surface methodology, are often sufficient for identifying major peaks, valleys, and ridges that define nutritional optima [81], outperforming generalized additive models and thin plate splines typically used to visualize response surfaces in GFN experiments [6, 82, 83, 84]. However, the adequacy of the model should always be evaluated through residual analysis, lack-of-fit tests, and cross-validation to ensure that the selected surface captures a genuine biological signal rather than experimental noise.

### 4.3 Multi-Objective Optimization in Mixture Experiments

Multi-objective optimization allows for the simultaneous maximization or minimization of multiple, often competing biological or performance objectives. In the context of nutritional experiments, this approach allows researchers to explore and balance trade-offs among several responses, such as growth, survival, reproduction, or cost of feed [55, 85]. When a statistical model, such as a polynomial regression, is fitted to experimental data, the predicted responses can be the subject of the optimization scheme. Two approaches which have been used in mixture experiments generally and nutritional experiments specifically are desirability functions [64, 86, 87] and Pareto optimization [88, 89].

Desirability functions are flexible tools that map each response onto a range from 0 to 1, where 1 represents the most desirable outcome [90]. Individual desirability scores (which correspond to different responses) can then be combined into a composite objective. When embedded within the framework of RSM, the object that gets optimized is not the raw experimental outcomes, but rather the prediction of the model fit to the data. In nutritional experiments, desirability functions can be meaningfully constrained by defining biologically relevant lower and upper bounds for each response variable. For example, growth rate may have a minimum acceptable value below which desirability is zero, while toxicity may have an upper bound beyond which desirability again falls to zero. Target-centered functions can also be used for responses with optimal intermediate values (i.e., body weight). The steepness and symmetry of these curves can be adjusted via shape parameters to reflect experimental priorities—for instance, penalizing underperformance more strongly than overconsumption. The overall composite desirability provides a scalar criterion for multi-response optimization (see Figure 8AB for an example).

**Figure 8:**
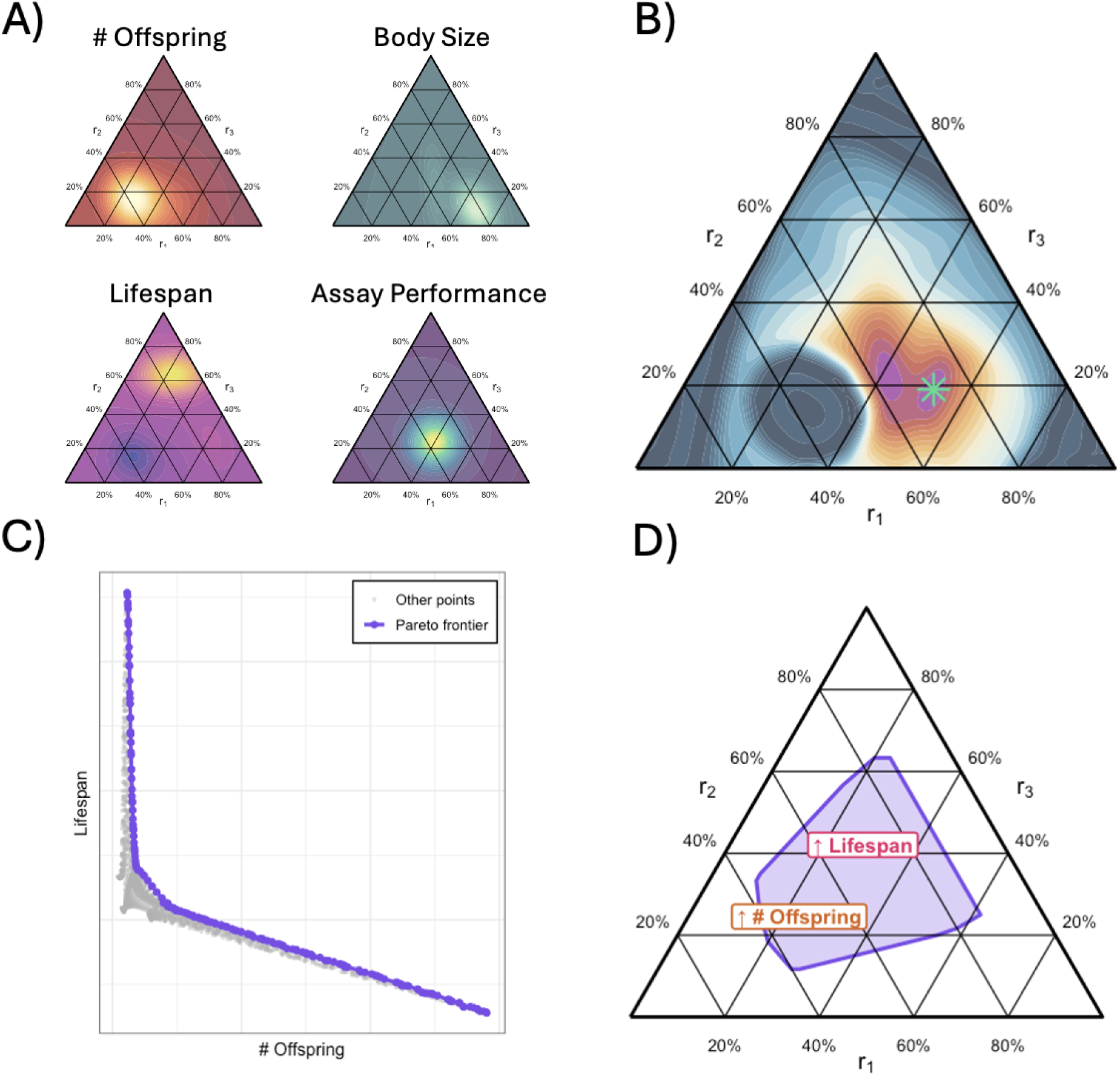
Multi-objective optimization in mixture experiments. A) Examples of response surfaces of various performance metrics from a nutrition experiment using the simplex. Data was generated from simulations and exists only for visualization purposes. Light colors indicate a maximum in each simplex. B) Response surface of the desirability function, which combines the responses from the first panel. The gold star indicates the global maximum. C) Scatterplot of the lifespan versus the number of offspring from these simulated experiments. The purple points are Pareto optimal. Points in the top right of the point cloud are considered optimal because the optimization goal for each variable is to maximize it. D) The purple region in the simplex corresponding to the previous panel. The left side of this polygon shows mixture combinations that maximize the number of offspring, whereas the top of this polygon shows where lifespan is maximized.

Pareto optimization represents an alternative but complementary approach to desirabilit based methods in mixture experiments. Rather than aggregating multiple objectives into a single scalar value, Pareto analysis preserves the individuality of each response, identifying a frontier of nondominated solutions—those for which no objective can be improved without worsening at least one other. This concept is particularly well suited to biological and nutritional systems, where the trade-off between competing physiological demands, such as reproduction versus longevity [55, 91] or growth versus immunity [92], should be analyzed fully. Within the GFN, Pareto optimization has been used to characterize such trade-offs in nutrient intake and performance across taxa [81, 60]. For instance, different combinations of macronutrients can yield optimal but distinct physiological outcomes, defining a continuous Pareto frontier of nutritional strategies that balance multiple aspects of fitness. By mapping these frontiers onto mixture coordinate systems, researchers can visualize the multidimensional structure of nutritional trade-offs and identify the regions of nutrient space where alternative strategies converge (Figure 8CD).

Desirability functions, Pareto frontiers, and other multi-objective approaches such as Bayesian or utility modeling are particularly relevant in animal growth and husbandry systems, where multiple measures of success coexist [6, 4, 89]. For example, the number of eggs, the number of survived group members, body temperature and mass, octopamine or cortisol levels, could all be indicators of individual health [25, 93]. In industrial settings such as insect farming, aquaculture, or cattle ranching, these methods can be used to design feed mixtures that maximize growth and efficiency while also maintaining animal welfare and minimizing cost or environmental impact. The same logic applies to laboratory settings, where multi-objective optimization can help standardize husbandry conditions for model organisms ranging from mice to social insects.

### 4.4 Case Study: Constrained Mixture Experiment for Nutritional Optimization

To illustrate the type of mixture experiments we recommend for nutritional studies, we constructed a simulated case study in which the goal was to identify the diet composition that maximizes either a single metric or a combination of metrics using something like desirability functions under some constraints: *r*_1_ *≤* 0.80, *r*_2_ *≤* 0.95, and *r*_3_ *≤* 0.70. These bounds carve out a feasible region inside the simplex and mimic the common situation in animal nutrition where one macronutrient (e.g. lipid) cannot be increased arbitrarily without violating palatability or welfare constraints. Here, the constraints should be set using a priori knowledge on what macronutrient contents will cause undue harm to the organism.

We first generated a dense candidate set of mixtures inside this constrained region and fit a quadratic Scheffé mixture model of the form

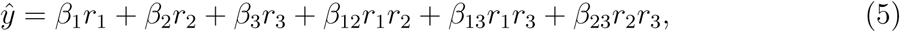

which is the standard second-order model for three-component mixtures. An A-efficient design was then selected from the candidate set using a row-exchange algorithm that minimized the trace of (**X**^⊤^**X**)^−1^, thereby producing a design that yields small average variances for the estimated coefficients. This step is critical in constrained mixture spaces because the usual simplex-lattice designs do not account for upper bounds on the ingredients and can place design points in infeasible portions of the ternary diagram. The selected design points were well distributed across the admissible region and included mixtures near the constraint boundaries, improving the ability of the fitted model to interpolate within the constrained simplex.

To emulate experimental data, we specified an underlying “true” response surface for growth whose maximum was deliberately placed near the interior of the feasible region. Each design point was then assigned a growth value from this surface with added Gaussian measurement error, representing variation among replicates or among individual organisms. Fitting the quadratic Scheffé model to these simulated data produced a smooth response surface over the constrained space. We then evaluated the fitted model on the full candidate set and identified the composition with the highest predicted growth; this served as the model-based optimum.

To assess how well the model-based optimum reflected observable performance, we ran five additional simulated replicates at the predicted optimal composition. For each of these replicates, the metric was drawn from the same underlying data-generating process used for the original design points. We then compared the model’s predicted metric at the optimum with the empirical distribution of these five replicates. In our simulation, the model-based prediction lay close to the replicate mean and within the range defined by the replicate standard deviation (predicted value = 53.3, the average replicate value = 51.9, RMSE = 1.59), indicating that the A-efficient design supported a well-conditioned fit of the mixture model, and the fitted surface was sufficiently accurate to guide follow-up experimentation. This workflow (constructing a constrained candidate set, choosing an optimal mixture design, fitting an appropriate Scheffé model, and validating the predicted optimum with targeted replication) provides a practical and ethical template for nutritional studies seeking to optimize growth under ingredient constraints (Figure 9).

**Figure 9:**
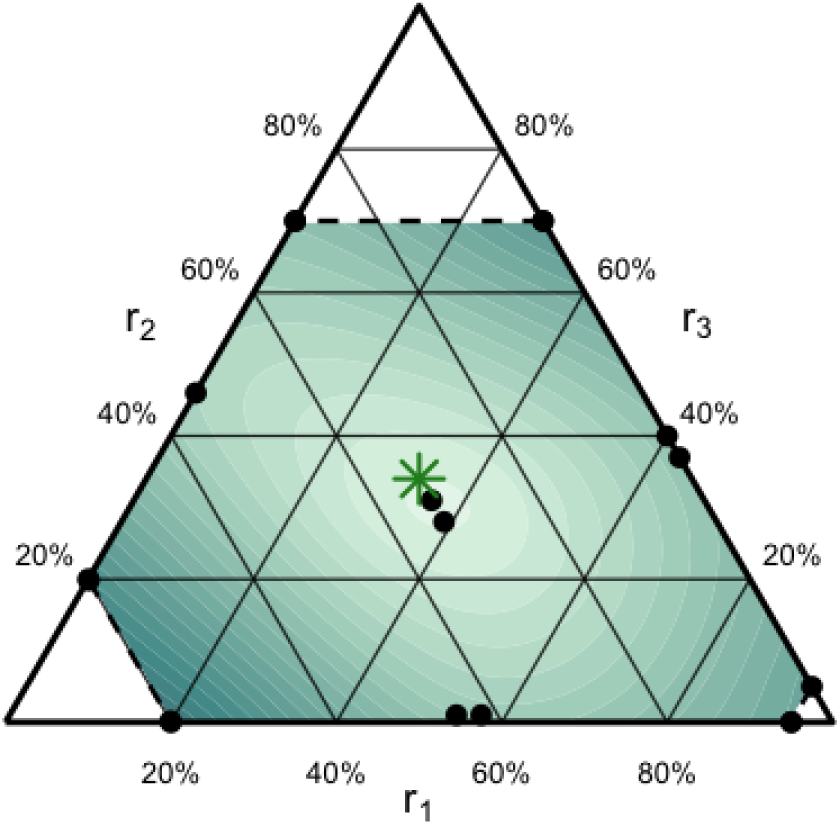
An example of a constrained experimental design which uses an optimality criterion to set design points. The response surface shows the value of the prediction of the quadratic Scheffé mixture model, which was fit to simulated data. The green star represents the highest point in this surface, and this is where 5 replicates were gathered to assess fit.

## 5. Discussion

In this paper, we have demonstrated how the simplex framework offers a coherent and statistically rigorous extension of the GFN by accommodating three-component mixtures in a balanced coordinate system. This contrasts with the RMT, which stretches the coordinate system along one of the axes and can thereby bias inferences. By situating concepts related to the GFN within the simplex, we are able to merge choice assays and no-choice (mixture) experiments under a common conceptual umbrella. This synthesis enables a more direct comparison of feeding behaviors (from choice experiments) and performance optima (from no-choice experiments), allowing growth targets derived from no-choice experiments to be meaningfully compared to intake targets from choice experiments and to random-choice baselines. Importantly, the statistical test for random choice introduced here gives practitioners a biologically interpretable null hypothesis in the mixture context: if the observed nutrient composition falls within the 95% confidence region of random sampling, then defense of a target ratio cannot be distinguished from stochastic feeding. This provides a more robust inferential foundation than many past studies, which have lacked a well-defined null model of food-choice behavior.

From an applied perspective, the implications are substantial [94, 95, 96]. In applied sectors, the use of optimal mixture-design principles and multi-objective optimization offers the possibility of identifying nutrient ratios that not only maximize growth or reproduction but also minimize cost, support sustainability goals, and improve welfare outcomes. Integrating health and welfare indicators into the objective space ensures that nutritional optimization in service to productivity or yield goals does not come at the expense of animal well-being, including in the insect farming context where animal welfare is increasingly recognized as an important part of precautionary, ethical practice [97, 98, 99, 100].

Constrained experimental designs also provide a clear ethical advantage in the research setting. By excluding nutritionally extreme or physiologically intolerable diet compositions, researchers avoid subjecting animals to conditions that are neither relevant in ecological or applied contexts nor necessary for generalizing study outcomes. This approach maximizes data integrity by ensuring that all sampled conditions fall within realistic, biologically meaningful boundaries, and it minimizes cost by focusing research effort on the most informative regions of nutrient space. In doing so, constrained mixture designs make more efficient use of public funding and reduces the environmental footprint associated with animal rearing and feed production. Thus, applying these principles to the design mixture experiments, researchers and practitioners can reduce sample sizes, limit potential harm, and derive more ethical and efficient feeding protocols.

The need for a structured nutritional framework becomes especially clear when considering recent studies that test large panels of diets without situating them within a common intellectual framework. For example, [85] evaluates the effects of approximately twenty different diets for yellow mealworms and reports substantial variation in multiple life-history traits, along with numerous apparent trade-offs. Yet, because the diets are not analyzed in terms of their relative positions in nutrient space, the biological drivers of these differences remain difficult to identify, no patterns relating life-history outcomes to macronutrients can be identified, and therefore the results cannot be readily generalized beyond the specific treatments used. This is a particular problem in insect farming contexts, where waste sources are often used; farmers may be constrained in how much they can do to make their diet ’match’ any particular diet used in a study because they lack choice in what substrate they can get access to. So, if a farmer only has bread bits, they cannot readily use information on wheat bran diets [85]. However, knowing the optimal P:F:C ratios would at least allow a rough mapping between available food sources and the ideal diet. A unified approach would also reduce waste at the scale of an entire field. For example, hundreds of choice assays have been performed on black soldier flies [101, 102, 103, 104], and yet a common consensus is difficult to reach due to a lack of a universal framework. A simplex-guided approach would permit the construction of response surfaces, the detection of nutrient trade-off boundaries, and a clearer understanding of how dietary composition shapes performance along many axes, allowing for a full synthesis of the effects of nutrition on model organisms.

From an experimental design standpoint, our reformulation outlines the importance of explicitly describing the statistical modeling pathway: the generation of a response surface in mixture space should not be merely glossed as “response-surface methodology”, but rather laid out in terms of model family (linear, GLM, mixture polynomial), transformation of compositional variables, validation diagnostics (residuals, lack-of-fit tests, cross-validation), and sensitivity to design choice. Without such rigor, the response surfaces may misrepresent true biological optima, and the potential for model bias or over-fitting remains. By combining rigorous mixture modeling with optimal design criteria that take real-world constraints into account (A, B, C, D-optimality), nutritional ecologists and animal nutrition scientists can ensure that their experimental efforts yield reliable estimates of nutrient-performance relationships while minimizing cost and reducing animal use. We specifically recommend the use of A-optimal designs in a constrained mixture space, where constraints are set by known limits on the macronutrient ratios of animals [105, 106, 107, 108].

Ultimately, by making three-component nutrition experimentally tractable and statistically robust using the simplex, our approach opens new pathways for both fundamental nutritional ecology and applied animal science, maximizes experimental efficiency and the ethical use of research animals, and improves decision-making for a variety of stakeholders working with animals in applied settings.

## Author Contributions

**Colin Lynch**: Conceptualization, Methodology, Software, Validation, Data curation, Writing—first draft, Writing—review. **Kaitlin Baudier**: Writing—review, Investigation, & editing. **Douglas Montgomery**: Writing—review, Investigation, & editing. **Meghan Barrett**: Writing—review, Investigation, & editing.

## Data Availability

The code used to generate figures will be made available upon reasonable request.

## Acknowledgments

We would like to thank Michaela Starkey for her comments on the manuscript.

