## supplementalMaterial for "Minimizing inferential bias in the theory and design of nutritional experiments through the application of the equilateral mixture triangle"

#### Supporting Information

#### S1: Defining the Simplex Coordinate System and Decision Rules

In this section, we formalize the simplex coordinate system for three components (Section S1.1), describe how the nutritional state of an organism evolves through sequential feeding decisions (Section S1.2), and introduce a set of probabilistic decision rules that map an organism's nutritional preferences to trajectories through this space (Section S1.3). This section ends with a conjecture as to which decision rule results in the optimal path to the intake target (Section S1.4). While the simplex can represent mixtures of any three interacting components, in nutritional ecology, it is most commonly used to represent macronutrient composition. Accordingly, we adopt the convention that the three mixture components $r_1$, $r_2$, and $r_3$ correspond to carbohydrate, protein, and fat, respectively. The same formalism, however, applies equally well to other mixtures such as amino acids, minerals, or energy sources.

##### S1.1: The Simplex Coordinate System

The EMT provides a natural coordinate system for representing animal nutritional states when three interacting macronutrients are considered. The EMT is mathematically equivalent to the simplex widely used in the design of experiments (DOE) for mixture problems [1, 2]. In this system, any food item is characterized by a three-dimensional composition vector of macronutrients (Figure 1), although in practice, each macronutrient could be replaced with other components of food:

$$\mathbf{F}_i = (F_i^{(c)}, F_i^{(p)}, F_i^{(f)}) \in \mathbb{R}^3, \tag{1}$$

where $F_i^{(c)}$, $F_i^{(p)}$, and $F_i^{(f)}$ are the proportions of carbohydrate, protein, and fat, respectively. These coordinates are constrained such that $F_i^{(c)} + F_i^{(p)} + F_i^{(f)} = 1$, meaning that every point lies on the 2-dimensional simplex defined by the convex hull of the three nutrient axes. Usually this simplex is plotted as an equilateral triangle, in which each vertex corresponds to a pure nutrient (100% of one macronutrient, 0% of the others), and any point inside the triangle corresponds to a feasible food or state of the organism (Figure 4). The current nutritional state of an organism at discrete time $t$ (which represents a distinct choice) is denoted as:

$$\mathbf{O}_t = (O_t^{(c)}, O_t^{(p)}, O_t^{(f)}), \qquad O_t^{(c)} + O_t^{(p)} + O_t^{(f)} = 1, \tag{2}$$

and the intake target is similarly represented by:

$$\mathbf{I} = (I^{(c)}, I^{(p)}, I^{(f)}), \qquad I^{(c)} + I^{(p)} + I^{(f)} = 1. \tag{3}$$

This coordinate system allows us to visualize trajectories of nutritional intake as an organism samples different food items.

#### S1.2: Evolution of the Nutritional State

Let $\mathcal{F} = \{\mathbf{F}_1, \ldots, \mathbf{F}_M\}$ denote the set of available foods, where each $\mathbf{F}_i = (F_i^{(c)}, F_i^{(p)}, F_i^{(f)})$ satisfies $F_i^{(c)} + F_i^{(p)} + F_i^{(f)} = 1$ and $F_i^{(\cdot)} \geq 0$. During feeding, the organism consumes nonnegative amounts $a_s \geq 0$ from foods indexed by $i_s \in \{1, \ldots, M\}$ across feeding bouts $s = 1, 2, \ldots, t$. The cumulative nutritional state of the organism after $t$ bouts is then the normalized mixture of all consumed foods:

$$\mathbf{O}_t = \frac{\displaystyle\sum_{s=1}^{t} a_s \, \mathbf{F}_{i_s}}{\displaystyle\sum_{s=1}^{t} a_s}. \tag{4}$$

Equivalently, let $A_i(t) = \sum_{s \leq t : i_s = i} a_s$ be the total amount consumed from food $i$ and $A(t) = \sum_{i=1}^{M} A_i(t)$ the overall intake. Then

$$\mathbf{O}_t = \sum_{i=1}^{M} \lambda_i(t) \, \mathbf{F}_i, \qquad \lambda_i(t) = \frac{A_i(t)}{A(t)}, \quad \lambda_i(t) \geq 0, \quad \sum_{i=1}^{M} \lambda_i(t) = 1. \tag{5}$$

Hence, at every point in time, the organism's nutritional state is a convex combination of the food compositions.

#### S1.3: Decision Rules in the Simplex Framework

We model animal foraging choices within the simplex by embedding nutritional states into a 3-dimensional Cartesian space constrained such that the sum of nutrient proportions equals one. Let the organism's nutritional state at time $t$ be denoted by $\mathbf{O}_t$, the intake target by $\mathbf{I}$, and the nutritional composition of food item $i$ by $\mathbf{F}_i$.

At each timestep, the organism evaluates the degree to which consuming food $i$ moves it toward $\mathbf{I}$. This is quantified by comparing the angle between the organism's current vector to the intake target and the vector from food $i$ to the intake target. Specifically,

$$\theta_{ti} = \cos^{-1}\left(\frac{(\mathbf{I}-\mathbf{O}_t)\cdot(\mathbf{I}-\mathbf{F}_i)}{\|\mathbf{I}-\mathbf{O}_t\|\,\|\mathbf{I}-\mathbf{F}_i\|}\right), \tag{6}$$

where $\theta_{ti}$ is the angular difference in radians between these two directional vectors.

The organism's propensity to select food $i$ is then mapped to a score $d_{ti}$ that peaks when $\theta_{ti} = \pi$ (i.e., the food vector is a reflection of the organism's trajectory toward the intake target):

$$d_{ti} = \cos(\theta_{ti} - \pi). \tag{7}$$

This function assigns higher values to food choices that align more closely with the desired trajectory toward $\mathbf{I}$.

Following a modified Boltzmann distribution, the probability of selecting food item $i$ is

$$P_{ti} = \frac{\exp(\kappa d_{ti})}{\sum_{j=1}^{M}\exp(\kappa d_{tj})}, \tag{8}$$

where $M$ is the number of available food items and $\kappa$ is an inverse temperature parameter controlling stochasticity. Low $\kappa$ yields near-random choice, while high $\kappa$ yields deterministic choice of the maximum $d_{ti}$.

Bias toward particular food items may be incorporated through weights $w_i$:

$$P_{ti} = \frac{w_i\exp(\kappa d_{ti})}{\sum_{j=1}^{M} w_j\exp(\kappa d_{tj})} \tag{9}$$

where $w_i$ sum to unity. Additionally, nutrient-specific preferences may be modeled by introducing a weight vector $\mathbf{W} = (w^{(c)}, w^{(p)}, w^{(f)})$ where again the weights sum to unity. The adjusted score is then

$$d'_{ti} = (\mathbf{W}\odot(\mathbf{I}-\mathbf{O}_t))\cdot(\mathbf{I}-\mathbf{F}_i), \tag{10}$$

where $\odot$ denotes elementwise multiplication. This formulation allows organisms to exhibit differential attraction to foods depending on their macronutrient content. The final decision rule is then given by:

$$P_{ti} = \frac{w_i \exp(\kappa d'_{ti})}{\sum_{j=1}^{M} w_j \exp(\kappa d'_{tj})}. \tag{11}$$

This decision-rule formalism provides a unified framework for modeling random foraging (low $\kappa$), deterministic optimization (high $\kappa$), food bias, and nutrient bias within the simplex. Importantly, the null expectation of random choice corresponds to convergence on the center of mass of the available food items, while deviations from this trajectory reflect systematic behavioral strategies.

Beyond their role as generic bias parameters, the food-specific weights $w_i$ can also be interpreted as quantities that map the EMT framework onto classical foraging theory. In particular, if each food item $i$ is associated with a net energetic payoff $E_i$ (for example, defined as the energetic gain per unit handling time) then setting $w_i \propto E_i$ effectively embeds optimal foraging considerations directly into the probabilistic choice rule. Foods yielding higher net energy gain, lower handling cost, or shorter search time naturally carry larger weights and therefore exert a stronger pull on the organism's trajectory through the simplex.

##### S1.4: Deterministic Optimality Conjecture

In the deterministic limit where $\kappa \to \infty$, with uniform food weights ($w_i = 1/M$) and equal nutrient weights $\mathbf{W} = (1/3, 1/3, 1/3)$, the probabilistic decision rule simplifies to a maximization principle. The organism deterministically selects the food item $i^*$ that maximizes the alignment score:

$$i^* = \arg \max_{i \in \{1,\ldots,M\}} d'_{ti}. \tag{12}$$

This yields a trajectory $\{\mathbf{O}_t^*\}_{t=1}^{T}$ that represents the most direct attainable path toward the intake target $\mathbf{I}$ given the available food set. Here, "most direct" is defined as the minimal number of steps required to reach $\mathbf{I}$ within a specified tolerance. We conjecture that, under this deterministic regime, the resulting trajectory is globally optimal in the sense that no other sequence of food choices can reach the same nutritional target in fewer steps without leaving the convex hull of available foods. This conjecture forms a theoretical benchmark against which stochastic or biased foraging strategies may be compared.

#### S2: Stretching in the RMT Coordinate Space

In the RMT, each composition $(r_1, r_2, r_3)$ satisfies the constraint $r_1 + r_2 + r_3 = 1$. Points in this coordinate system are projected onto a two-dimensional Cartesian plane according to the mapping

$$(r_1, r_2, r_3) \;\mapsto\; (r_1, r_2), \tag{13}$$

so that the proportion of the third component is given implicitly by $r_3 = 1 - r_1 - r_2$. This transformation defines an equilateral coordinate system in which small displacements along the $r_1$ and $r_2$ axes correspond to equal Euclidean steps in the underlying space, whereas motion along the $r_3$ direction produces a larger displacement.

To determine the relative scaling of these directions, consider infinitesimal changes in each coordinate that preserve the compositional constraint $r_1 + r_2 + r_3 = 1$. A displacement along the $r_1$ direction by an amount $\Delta r_1$ maps to $(\Delta r_1, 0)$ in the projected plane, corresponding to a Euclidean distance $\|\Delta\mathbf{r}\| = |\Delta r_1|$. Similarly, a displacement along the $r_2$ direction by $\Delta r_2$ maps to $(0, \Delta r_2)$, producing a distance $\|\Delta\mathbf{r}\| = |\Delta r_2|$.

In contrast, a change in the third component $r_3$ must be accompanied by adjustments in $r_1$ and $r_2$ to maintain the constraint. Because $r_1 + r_2 + r_3 = 1$, a small increase $\Delta r_3$ implies $\Delta r_1 = \Delta r_2 = -\frac{1}{2}\Delta r_3$. The corresponding displacement in the plotted plane is then $\big(-\frac{1}{2}\Delta r_3, -\frac{1}{2}\Delta r_3\big)$, whose Euclidean magnitude is

$$\|\Delta\mathbf{r}\| = \sqrt{\left(\tfrac{\Delta r_3}{2}\right)^2 + \left(\tfrac{\Delta r_3}{2}\right)^2} = \frac{|\Delta r_3|}{\sqrt{2}}. \tag{14}$$

Hence, a unit change in $r_3$ corresponds to a displacement of $1/\sqrt{2}$ in the projected plane, whereas unit changes in $r_1$ or $r_2$ corresponds to unit displacements. To produce the same Euclidean step length as a unit change in $r_1$ or $r_2$, the $r_3$ axis must therefore be scaled by a factor of $\sqrt{2}$. Equivalently, motion along the $r_3$ direction in the RMT coordinate system can be viewed as being "stretched" by $\sqrt{2}$ relative to motion along the $r_1$ or $r_2$ axes.

### S3: Convex Hull Contains All Possible Outcomes

**Proposition.** Let $\mathcal{F} = \{\mathbf{F}_1, \ldots, \mathbf{F}_M\} \subset \Delta_2$ be the set of available food compositions satisfying $F_i^{(c)} + F_i^{(p)} + F_i^{(f)} = 1$. If the organism's nutritional state $\mathbf{O}_t$ evolves according to Eq. 4, then for all $t \geq 1$,

$$\mathbf{O}_t \in \mathrm{conv}(\mathcal{F}), \tag{15}$$

where $\mathrm{conv}(\mathcal{F})$ denotes the convex hull of the food set. Conversely, every point of $\mathrm{conv}(\mathcal{F})$ can be approximated arbitrarily closely by some sequence of consumptions $\{a_s, i_s\}$.

**Proof.** Let $A_i(t)$ and $A(t)$ be defined as above. Then by Eq. 5, $\mathbf{O}_t = \sum_{i=1}^{M} \lambda_i(t)\mathbf{F}_i$ with $\lambda_i(t) \geq 0$ and $\sum_i \lambda_i(t) = 1$, which by definition means $\mathbf{O}_t \in \mathrm{conv}(\mathcal{F})$. For the converse, take any $\mathbf{x} = \sum_{i=1}^{M} \lambda_i \mathbf{F}_i$ with $\lambda_i \geq 0$ and $\sum_i \lambda_i = 1$. Choose feeding sequences such that the empirical proportions $A_i(t)/A(t) \to \lambda_i$. Then by continuity of Eq. 5, $\mathbf{O}_t \to \mathbf{x}$. Hence the convex hull of the available foods exactly bounds the space of all attainable nutritional outcomes.

### S4: Mathematical Derivation of the Confidence Ellipse for Random Food Choice

Let $\mathbf{r}_j = (r_{1j}, r_{2j}, r_{3j})^\top$ denote the barycentric coordinates of the $j^{\text{th}}$ food item, where $r_{1j}, r_{2j}, r_{3j} \geq 0$ and $r_{1j}+r_{2j}+r_{3j} = 1$. These coordinates represent the relative proportions of three nutrients (for example, carbohydrate, protein, and fat) in food $j$, and the index $j \in \{1, 2, \ldots, M\}$ enumerates the $M$ available foods. Under the assumption of random choice, all foods are selected with equal probability $p_j = 1/M$. The expected nutrient ratio of a randomly selected food is given by

$$\bar{\mathbf{r}} = \sum_{j=1}^{M} p_j \, \mathbf{r}_j, \tag{16}$$

which represents the center of mass of the available foods (plotted as the blue point in Figure 4). The variability in random draws from this uniform distribution can be described by the population covariance matrix

$$\Sigma_r = \sum_{j=1}^{M} p_j \, (\mathbf{r}_j - \bar{\mathbf{r}})(\mathbf{r}_j - \bar{\mathbf{r}})^\top. \tag{17}$$

Because the mixture space is constrained to $r_1 + r_2 + r_3 = 1$, this covariance matrix has rank two. To visualize these relationships in the simplex, the barycentric coordinates are projected into two-dimensional Cartesian coordinates,

$$\mathbf{x}_j = \begin{pmatrix} x_j \\ y_j \end{pmatrix} = \begin{pmatrix} r_{2j} + \frac{1}{2} r_{3j} \\ \frac{\sqrt{3}}{2} \, r_{3j} \end{pmatrix}, \tag{18}$$

so that each food occupies a unique position within the triangular space. The corresponding two-dimensional covariance matrix is then

$$\Sigma_{xy} = \sum_{j=1}^{M} p_j \, (\mathbf{x}_j - \bar{\mathbf{x}})(\mathbf{x}_j - \bar{\mathbf{x}})^\top, \qquad \bar{\mathbf{x}} = \sum_{j=1}^{M} p_j \, \mathbf{x}_j. \tag{19}$$

If the mean nutrient ratio is estimated from $n$ independent choices drawn from the $M$ foods, the sampling covariance of the estimated mean is

$$\Sigma_{\bar{x}y} = \frac{\Sigma_{xy}}{n}. \tag{20}$$

Assuming that the sample mean is approximately bivariate normal,

$$\bar{\mathbf{x}}_{\text{sample}} \sim \mathcal{N}_2(\bar{\mathbf{x}}, \Sigma_{\bar{x}y}), \tag{21}$$

the $(1-\alpha)$ confidence region for $\bar{\mathbf{x}}$ is the interior of the ellipse satisfying

$$(\mathbf{x}-\bar{\mathbf{x}})^\top \Sigma_{\bar{x}y}^{-1}(\mathbf{x}-\bar{\mathbf{x}}) \leq \chi^2_{2,1-\alpha}, \tag{22}$$

where $\chi^2_{2,1-\alpha}$ is the quantile $(1-\alpha)$ of the chi-square distribution with two degrees of freedom. If we let $\Sigma_{\bar{x}y} = Q\Lambda Q^\top$ denote the eigendecomposition of the sampling covariance matrix, with $\Lambda = \mathrm{diag}(\lambda_1, \lambda_2)$ and $Q$ an orthonormal rotation matrix, then the ellipse can be parameterized as

$$\mathbf{x}(\theta) = \bar{\mathbf{x}} + \sqrt{\chi^2_{2,1-\alpha}}\, Q\Lambda^{1/2} \begin{pmatrix} \cos\theta \\ \sin\theta \end{pmatrix}, \qquad \theta \in [0, 2\pi). \tag{23}$$

The semi-axis lengths of the ellipse are therefore

$$a = \sqrt{\chi^2_{2,1-\alpha}\,\lambda_1}, \qquad b = \sqrt{\chi^2_{2,1-\alpha}\,\lambda_2}, \tag{24}$$

with orientation given by the principal eigenvector of $\Sigma_{\bar{x}y}$. The resulting ellipse encloses the region of the simplex in which 95% of sample means from random food choice would be expected to occur. If an observed intake target or feeding trajectory falls outside this region, it indicates a significant deviation from random expectation, implying directed regulation or preference in nutrient selection. The mathematical derivation of this confidence region follows standard results from multivariate normal theory (see [3]).

### S5: The Design Matrix in Response Surface Methodology

In response surface methodology (RSM), the design matrix $\mathbf{X}$ encodes the structure of the experimental design and forms the foundation for model estimation. Each row of $\mathbf{X}$ corresponds to a single experimental run, while each column represents one of the model terms—typically including a constant (intercept), linear effects, interaction terms, and higher-order polynomial components depending on the complexity of the fitted model.

For example, in a second-order (quadratic) model with $k$ factors, the design matrix is expressed as

$$\mathbf{X} = \begin{bmatrix} 1 & x_{11} & x_{12} & \cdots & x_{1k} & x_{11}^2 & x_{11}x_{12} & \cdots & x_{1k}^2 \\ 1 & x_{21} & x_{22} & \cdots & x_{2k} & x_{21}^2 & x_{21}x_{22} & \cdots & x_{2k}^2 \\ \vdots & \vdots & \vdots & & \vdots & \vdots & \vdots & & \vdots \\ 1 & x_{n1} & x_{n2} & \cdots & x_{nk} & x_{n1}^2 & x_{n1}x_{n2} & \cdots & x_{nk}^2 \end{bmatrix},$$

where $n$ is the total number of experimental runs and $x_{ij}$ denotes the coded level of factor $j$ in run $i$ (as opposed to the empirical unit). The corresponding response vector $\mathbf{y}$ is related to $\mathbf{X}$ through the standard linear model formulation

$$\mathbf{y} = \mathbf{X}\boldsymbol{\beta} + \boldsymbol{\varepsilon}, \tag{25}$$

where $\boldsymbol{\beta}$ is the vector of unknown regression coefficients and $\boldsymbol{\varepsilon}$ represents random experimental error.

Thus, $\mathbf{X}$ serves as the mathematical link between the experimental design and the fitted response surface, determining both the estimability and precision of the model coefficients. The choice of design (reflected in the structure of $\mathbf{X}$) directly influences the interpretability of the model, the precision of the prediction, and the ability to detect curvature or interactions in the response.

### S6: Assessing Rules of Compromise via Trajectory Matching

The probabilistic decision framework introduced in Section S1.3 provides a general means of simulating how organisms move through the nutrient ratio space of the simplex. By adjusting its parameters, one can generate idealized trajectories that correspond to specific rules of compromise, which are heuristics describing how organisms regulate nutrient intake when faced with suboptimal or imbalanced food choices. These theoretical trajectories provide reference paths against which observed behavior can be quantitatively compared.

An experimental trajectory records how the organism's nutritional state changes over time:

$$\mathbf{O} = \{\mathbf{O}_t\}_{t=1}^{T},$$

where $\mathbf{O}_t$ is the organism's state in simplex coordinates after the $t^{\text{th}}$ feeding bout. Each theoretical trajectory,

$$\mathbf{O}' = \{\mathbf{O}'_t\}_{t=1}^{T'},$$

represents an idealized rule of compromise, generated from the decision rule in Section S1.3 under particular limiting cases or parameterizations.

The deterministic optimal trajectory ($\mathbf{O}'_{\text{opt}} = \{\mathbf{O}^*_t\}_{t=1}^{T}$) is obtained by taking the limit $\beta \to \infty$ with equal food and nutrient weights. In this case, the organism always selects the food that most directly advances its state toward the intake target, producing the shortest feasible path through nutrient space. This corresponds to the "closest distance optimization" rule of compromise.

The single-food trajectory ($\mathbf{O}'_{\text{food}}$) results from assigning weight $w_i = 1$ to one food and $w_j = 0$ to all others. In the deterministic limit, the organism consumes only that food, and its state converges to the food's composition. This represents exclusive preference for a single food item.

The macronutrient-prioritization trajectory ($\mathbf{O}'_{\text{mac}}$) is generated by assigning unequal nutrient weights $\mathbf{W} = (w^{(c)}, w^{(p)}, w^{(f)})$ such that one nutrient dominates (e.g., $w^{(c)} \gg w^{(p)}, w^{(f)}$). The organism first reduces its deficit in that nutrient before adjusting others as weights are relaxed toward equality. This captures the sequential nutrient-balancing strategies described in amount-based geometric framework studies.

Finally, the random-choice trajectory ($\mathbf{O}'_{\text{rand}}$) is obtained by setting all weights equal and taking the stochastic limit $\beta \to 0$. All foods are sampled uniformly, and the expected path converges on the center of mass of the convex hull of available foods (see Section S4). This trajectory serves as the null model of unregulated choice.

To quantify how closely an observed trajectory $\mathbf{O}$ follows a theoretical rule $\mathbf{O}'$, we compute a distance between their paths in simplex space. When the trajectories have comparable time resolution, a simple lockstep Euclidean distance may suffice:

$$D_{\text{L}}(\mathbf{O}, \mathbf{O}') = \sum_{t=1}^{\min(T,T')} \|\mathbf{O}_t - \mathbf{O}'_t\|. \tag{26}$$

However, when the organism moves through nutrient space at a different rate than the theoretical trajectory, a dynamic time warping (DTW) distance is preferred:

$$D_{\text{W}}(\mathbf{O}, \mathbf{O}') = \min_{\pi} \sum_{(t,t')\in\pi} \|\mathbf{O}_t - \mathbf{O}'_{t'}\|, \tag{27}$$

where $\pi$ denotes a monotonic warping path that aligns points between the two sequences. DTW allows trajectories to be compared based on shape rather than strict temporal correspondence.

Each distance $D(\mathbf{O}, \mathbf{O}')$ can be converted into a similarity score using the exponential decay function,

$$S = \exp[-\alpha D(\mathbf{O}, \mathbf{O}')], \qquad \alpha > 0, \tag{28}$$

and normalized across a set of candidate rules $\{\mathbf{O}'_k\}$ as

$$\tilde{S}_k = \frac{S_k}{\sum_j S_j}. \tag{29}$$

The normalized scores $\tilde{S}_k$ represent the relative support for each hypothesized rule of compromise.

This approach allows empirical feeding trajectories to be evaluated against theoretical behavioral rules in a unified quantitative framework, identifying whether observed foraging patterns are best explained by random choice, single-food attraction, nutrient prioritization, or directed optimization toward the intake target.
